# A universal RT-qPCR assay for “One Health” detection of influenza A viruses

**DOI:** 10.1101/2020.06.29.171306

**Authors:** Alexander Nagy, Lenka Černíková, Kateřina Kunteová, Zuzana Dirbáková, Saumya S Thomas, Marek J Slomka, Ádám Dán, Tünde Varga, Martina Máté, Helena Jiřincová, Ian H Brown

**Author notes:** **Emails:** LČ, KK, ZD, SST, MJS, ÁD, TV, MM, HJ, IHB. **Corresponding author** Alexander Nagy, State Veterinary Institute Prague, Sídlištní 136/24, 165 03 Prague 6, Czech Republic.

## Abstract

The mutual dependence of human and animal health is central to the One Health initiative as an integrated strategy for infectious disease control and management. A crucial element of the One Health includes preparation and response to influenza A virus (IAV) threats at the human-animal interface. The IAVs are characterized by extensive genetic variability, they circulate among different hosts and establish host-specific lineages. The four main host reservoirs are: avian, swine, human and equine, with occasional transmission to other mammalian species. The host diversity is mirrored in the range of the RT-qPCR assays for IAV detection. Different assays are recommended by the responsible health authorities for generic IAV detection in birds, swine or humans. In order to unify IAV monitoring in different hosts and apply the One Health approach, we developed a single RT-qPCR assay for universal detection of all IAVs of all subtypes, species origin and global distribution. The assay design was centred on a highly conserved region of the IAV MP-segment identified by a comprehensive analysis of 99,353 sequences. The reaction parameters were effectively optimised with efficiency of 93-97% and LOD_95%_ of approximately ten IAV templates per reaction. The assay showed high repeatability, reproducibility and robustness. The extensive *in silico* evaluation demonstrated high inclusivity, i.e. perfect sequence match in the primers and probe binding regions, established as 94.6% for swine, 98.2% for avian and 100% for human H3N2, pandemic H1N1, as well as other IAV strains, resulting in an overall predicted detection rate of 99% on the analysed dataset. The theoretical predictions were confirmed and extensively validated by collaboration between six veterinary or human diagnostic laboratories on a total of 1970 specimens, of which 1455 were clinical and included a diverse panel of IAV strains.

## Introduction

In 2004 the World Health Organization (WHO), the World Organization for Animal Health (OIE), the United Nations’ Food and Agriculture Organization and the World Bank proposed a One Health Initiative as a holistic perspective on the management and control of influenza A virus (IAV) threats at the human-animal interface [1]. The One Health concept is based on the realization that human and animal health allied to environmental safeguarding are inextricably linked [2]. All IAVs originate from the wild bird reservoir across a very wide geographical range. They include many different genotype combinations of the eight genomic segments, but all IAVs are classically subdivided to subtypes defined by the hemagglutinin (HA) and neuraminidase (NA) glycoprotein combinations. The mechanism of variability includes drift which affects all eight genetic segments, with additional genetic diversity and scope for viral evolution provided by reassortment among the segments. In the case of HA and NA gene segments, their reassortment (genetic and associated antigenic shift) has featured in the emergence of novel human IAV pandemics [3, 4].

Along with the established human seasonal IAVs [5, 6], important animal IAVs include the avian influenza viruses (AIVs) [7–9] and swine influenza viruses (SwIVs) [10, 11]. AIVs in particular are recognized as a key potential threat to poultry and human health globally, with the notifiable H5 and H7 AIV subtypes having the potential to mutate from low pathogenicity (LP)AIV to the corresponding highly-pathogenic (HP)AIV with consequent virulent infections in farmed birds [7, 9]. Pigs are another important host which has been long known to serve as a “mixing vessel” for IAVs of different species origins, enabling genetic reassortment to generate new circulating SwIV strains in pigs which may potentially transmit zoonotically [12, 13]. Therefore, collaboration between human and animal health experts is prudent to respond to the threats posed by different IAV strains, particularly where a newly-evolved strain may be threatening to cross or has crossed the species barrier, and harmonise the appropriate prevention and control steps [14]. The One Health approach prepared for the threat of goose/Guangdong (GsGd) lineage H5N1 HPAIVs [9] to both poultry and public health, and also to the 2009 pandemic H1N1 IAV (H1N1pdm09) which spread globally in humans [15] and also became established in pigs [11].

A key component of the One Health concept is to ensure our ability to sensitively, specifically and universally detect the IAVs from different host species and sample matrices. The relatively conserved bicistronic genomic segment no. 7 (matrix protein (MP)-segment) of the viral genome is an attractive region for generic IAV detection by RT-qPCR. Primer and probe binding regions were previously selected within the most conserved regions of the AIV MP-segment, to guide the design and validation of an RT-qPCR for the global detection of AIVs belonging to all sixteen HA-subtypes [16]. Blind trials in a consortium of five European AI reference laboratories [17] led to international acceptance of this generic AIV detection protocol by the European Union and the OIE [18–20]. This assay has proven itself as a sensitive test in the initial diagnosis of suspect AIV outbreaks, prompting subsequent investigations of notifiable AIV by means of type-specific H5 and H7 RT-qPCRs [21, 22]. However, continuing genetic drift within the AIV MP-segment of newly-emerging clades of the GsGd lineage epizootic H5Nx HPAIVs compromised assay sensitivity due to nucleotide mismatches in the reverse primer binding sequence [23]. In addition, a more recently designed RT-qPCR which amplifies elsewhere within the MP-segment was validated [24] and featured successfully in subsequent AIV outbreaks and wild bird investigations [25–28], AIV *in vivo* studies [29, 30] and SwIV field investigations [31–34]. The importance of appropriate test validation has been emphasised in a veterinary context [35], with an MP-segment RT-qPCR for generic detection of contemporary H1N1, H1N2 and H3N2 SwIV subtypes in European pigs having been also validated [36].

In the case of human seasonal IAVs, a recent WHO protocol lists four different MP-segment RT-qPCR assays to encompass both seasonal and potential zoonotic strains [37]. Three MP RT-qPCRs continue to be listed by the OIE for IAV detection in swine, horses and birds [38–40]. Many of these assays were designed and validated in the context of IAV detection in a single species group. Although the MP-segment is relatively conserved in the IAV genome, certainly when compared to the subtype-determining HA and NA genes, continuing genetic drift has contributed to considerable nucleotide variability [41–43]. Therefore, there is an ongoing risk that the unpredictable emergence of new IAV strains with critical mismatches in the primer and/or probe binding sequences may compromise the sensitivity of the RT-qPCR assay, therefore requiring assay modifications or replacement [23, 36, 44, 45]. Such primer or probe complementarity problems may only be recognised retrospectively, when outbreaks, epizootics or epidemics are already ongoing. In addition, the history of human IAV pandemics has underlined the importance of retaining the ability to detect historical, re-emerging IAV strains [46, 47]. In the case of H1N1pdm09, its MP-segment had apparently circulated unrecognized in swine for roughly for a decade [15]. Given these challenges, consideration of a single generic approach for universal IAV detection of all subtypes remains highly desirable, particularly in the relevant One Health species groups. This study describes the careful modification of the MP-segment RT-qPCR [24] based on the analyses of 99,353 of IAV MP-segment sequences available in the public sequence databases, together with its validation to fulfil this test criterion. Crucially, the modified assay design employed novel, more refined, *in silico* bioinformatics approaches [43].

## Materials and methods

### Clinical specimens, viral and bacterial taxa

Three specimen groups were included in the assay validation: i) 949 IAV negative field specimens collected from six mammalian and 68 avian species, ii) 409 non-IAV viruses, bacteria, and vaccine strains iii) 612 IAV positive specimens. The IAV positive pool was composed of 270 AIVs in 49 subtype combinations (both HPAIV and LPAIV, all HA and NA subtypes, collected from 1956 to 2020), 120 swine H1N1, H1N2 and H3N2 viruses (collected from 1946 to 2019), 108 human H3N2, 66 human pandemic H1N1 (H1N1pdm), 18 human pre-2009 and eight zoonotic strains (collected from 1934 to 2019) and 21 equine IAVs (collected from 1956 to 2009). The IAV positive pool included 369 IAV laboratory isolates and 243 characterised IAV positive field specimens. The field specimens encompassed various sample matrices like cloacal, tracheal and nasopharyngeal swabs, faeces, pooled organ suspensions and bronchoalveolar lavages. The list of all specimen included in the study was provided in Supporting information 1.

All specimens were obtained from the repositories of the collaborating institutions. All of the sensitive data were treated anonymously according to the international standard EN ISO/IEC 17025: 2018 and the entire data management is compliant with General Data Protection Regulation of the European Union.

### Quantified standard (QS)

Total nucleic acid extracts of A/mallard/Czech Republic/13579-84K/2010(H4N6) and A/goose/Czech Republic/1848-T14/2009(H7N9) AIVs (further referred as QS-1 and QS-2 respectively; GenBank accession numbers <u>JF789621</u> and <u>HQ244418</u>) were quantified by comparison to a dilution series of a known quantity of synthetic SVIP-MP amplicon (4nM DNA ultramer, Integrated DNA Technologies) tested by the SVIP-MP assay version 1 [24]. The ultramer dilution series spanned a concentration range from 1e7 to 1e2 copies/μl in triplicate. The standards were aliquoted and stored at −80°C.

### Nucleic acid extraction

Nucleic acid extraction was performed by accredited means within the collaborating laboratories: i) MagNA Pure Compact (Total Nucleic Acid Extraction Kit, input volume of 200 or 400μl and an elution volume of 50μl) and MagNA Pure 96 (DNA and Viral NA Small Volume Kit, input volume of 200 and an elution volume of 50μl) extractors (all from Roche), ii) Nucleomag Vet kit (Macherey-Nagel) or MagAttract 96 cador Pathogen Kit (Qiagen) on King Fisher Flex 96 (Thermo Fisher Scientific): Input volume 200 μl and elution volume 100μl, and iii) manual and robotic extraction depending on the origin of the specimen, as described by Slomka et al. [48].

### Primers and probe selection

The primers and the probe (Table 1) were selected upon a comprehensive *in-silico* analysis of 99,353 AIV MP sequences which represented the entire collection of the IVD (https://www.ncbi.nlm.nih.gov/genomes/FLU/Database/nph-select.cgi?go=database) and GISAID’s EpiFlu (https://www.gisaid.org/) databases collection as of February 2018 [43]. Briefly, the sequences were downloaded in five data pools (Table 2) and aligned with the MAFFT tool [49]. Alignment processing was performed with the AliView program [50]. Positional nucleotide numerical summary (PNNS) and entropy were calculated by the PNNS calculator and Entropy Calculator modules of the Alignment Explorer web application [51].

**Table 1.**
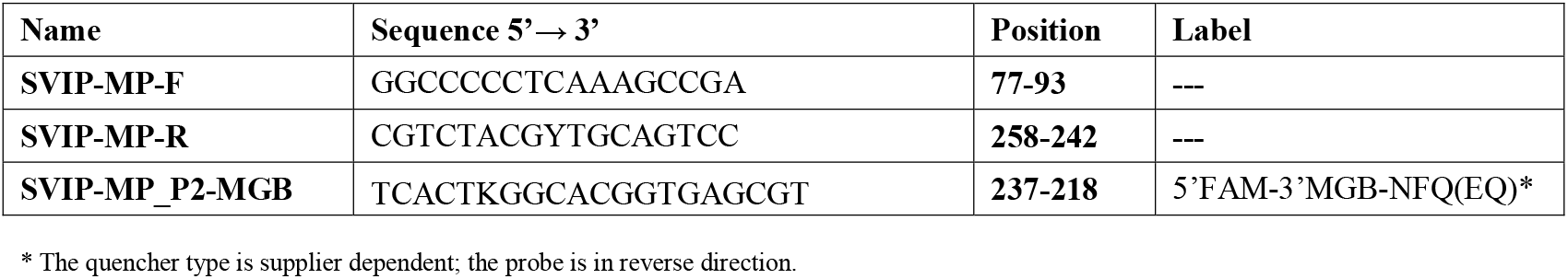
The primer and probe details.

**Table 2.** The MP-segment data groups, database origin and SVIP-MPv2 assay theoretical inclusivity values.

| Data groups |  | Database source* | Downloaded MP-sequences | Number of informative sequences<sup>#</sup> | Assay inclusivity |
| --- | --- | --- | --- | --- | --- |
| Human H3N2 |  | GISAID | 37,406 | 35,876 | 35,876 (100%) |
| Human H1N1pdm |  | GISAID | 16,948 | 16,311 | 16,311 (100%) |
| Avian |  | GISAID | 21,530 | 20,652 | 20,287 (98.2%) |
| Swine |  | IVD | 9,337 | 9,088 | 8,596 (94.6%) |
| Others | Others subtotal |  | 7,064 | 5,927 | 5,927 (100%) |
|  | Human H1N1 seasonal | IVD | 3,140 |  |  |
|  | Human H7Nx | GISAID | 1,249 |  |  |
|  | Human H5Nx | GISAID | 428 |  |  |
|  | Human H2N2 | IVD | 129 |  |  |
|  | Human H1N2 | GISAID | 71 |  |  |
|  | Human H9N2 | GISAID | 22 |  |  |
|  | Human H10N8 | IVD | 6 |  |  |
|  | Environment | GISAID | 1,368 |  |  |
|  | Canine | IVD | 303 |  |  |
|  | Equine | GISAID | 238 |  |  |
|  | Feline | GISAID | 44 |  |  |
|  | Other mammals | GISAID | 70 |  |  |
| **TOTAL** |  |  | **99,353** | **87,854** | **86,997 (99.0%)** |

\* GISAID- EpiFlu database, Global Initiative on Sharing All Influenza Data; IVD-Influenza Virus Database, National Centre for Biotechnology Information [52]. Sequences downloaded as of February 2018.
<sup>#</sup> MP-segment submissions containing the primer and probe sequences.

### Inclusivity assessment *in-silico*

The aligned MP-segment data pools were trimmed and concatenated (joining only the primer-probe-primer binding motifs into a continuous sequence; further referred as region of interest, ROI; Supporting dataset) with the AliView program [50]. Alignment stratification and sequence sorting was performed by the SequenceTracer module of the Alignment Explorer web application. The sequence variants in the alignment were considered at above the 0.5% threshold level. For a detailed analysis pipeline please refer to [43]. For additional sequence sorting according to isolate names the Fasta headers were extracted with FaBox 1.42 http://users-birc.au.dk/palle/php/fabox/ [53].

### SVIP-MPv2 RT-qPCR assay

The SVIP-MP assay version 2 (State Veterinary Institute Prague) consists of the original SVIP-MP-F and SVIP-MP-R primer pair [24] (Integrated DNA Technologies), and a newly designed 5’ FAM-labelled and 3’-minor groove binder (MGB)-modified probe SVIP-MP_P2-MGB ((Table 1; ThermoFisher Scientific, Eurofins Genomics). The primers encompass a 182 nucleotides-long amplicon. The probe is synthesised in the reverse complement orientation, i.e. its 5’-end is digested downstream from the reverse primer. The SVIP-MPv2 assay was developed and validated as a one-step procedure using the QuantiNova Probe RT-PCR Kit (Qiagen) with final concentrations of primers (1.4μM each) and probe (0.4μM) a final volume of 25μl (20μl reaction mix and 5μl of total NA extract).

The reactions were run on the CFX 96 (BioRad) thermal cycler with the following thermoprofile: 45°C 10min, 95°C 5min followed by 45 cycles of 95°C for 10s and 64°C for 30s and 72°C 10s, with signal acquisition in the FAM channel at the end of the 64°C annealing step. The Cq values were estimated by using the automatic baseline threshold option of the CFX Manager software v3.1 or CFX Maestro software v1.1.

The diagnostic test criteria of the SVIP-MPv2 assay were evaluated as described previously [54–56] with general performance criteria defined according to [55].

MeltMan [57] SVIP-MPv2 RT-qPCR reactions were performed by adding the SYTO82 dye (Thermo Fisher Scientific) in a final concentration of 0.8μM, without changing other reaction components, and the thermoprofile was extended with melting analysis from 50°C to 95°C and continuous signal gathering in the VIC channel of the CFX96 instrument.

### Efficiency

Calibration and standard curves were constructed by 10-log dilution of the QS-1 and QS-2 in a concentration from 1e6 to 1e2 (QS1) or from 1e7 to 1e2 (QS-2) copies/μl in triplicates. The dilutions were prepared in a total background nucleic acid of swine origin (at least 7ng/μl) and analysed on the CFX96 instrument. In addition, the efficiency was inferred by using linear regression analysis in the LinReg v. 2017.1 program [58].

### Limit of Detection (LOD)

The LOD was estimated according to the modified Two LOD procedure [55] and expressed as IA virions/μl of NA extract. First, the LOD_6_ was determined by preparing a serial dilution of the QS in a range of 100, 50, 20, 10, 5, 2, 1 and 0.1 copies/μl in six replicates and two independent runs resulting in a total of 12 replicates per each dilution point. All dilutions series were prepared in a total swine genomic nucleic acid background with a concentration at least 7ng/μl under repeatability conditions (i.e. prepared separately before each run). The highest QC dilution with all replicates positive was considered as the LOD_6_ of the assay. The accuracy of the dilution series was verified at the 0.1 dilution level where a maximum of one positive result per six replicates was accepted. The LOD of the assay with 95% confidence level (LOD_95%_) was determined by 60 replicates of the QS dilution corresponding to the LOD_6_ on the condition that all 60 replicates must have shown specific positive amplification. Finally, the LOD_95%_ was calculated by constructing the Probability of Detection (POD) curve [56].

### Precision

The precision is related to the random error of measurement and characterizes the closeness of the measured values obtained by replicate measurements under specified conditions. The precision is multicomponential. We considered the repeatability (intra-assay variation) and reproducibility (inter-assay variation). The repeatability and reproducibility of the SVIP-MPv2 assay was estimated from three measurements of 12 replicates of the QS-1 in the concentration of 10x LOD_95%_ prepared in a total swine genomic NA background (at least 7ng/μl). Each measurement was performed on different days using different CFX96 instruments where the reaction mixes were prepared by three different operators using different pipette sets. The repeatability was calculated for each run and was expressed as standard deviation (SD) and repeatability relative standard deviation as a percentage (%RSD_r_). The reproducibility was investigated from all three runs and the results were expressed as reproducibility relative standard deviation as a percentage %RSD_R_ [55].

### Robustness

The robustness assesses the capacity of the assay to tolerate small deviations during the experiment preparation and measurement [56]. It can also reveal instrument or qPCR kit dependence. The assay robustness was determined by implementing the fractional factorial experimental design [55, 56]. A six-parameter orthogonal combination matrix was designed to follow multiple parameters at any given time (Supporting information 2, Table S1). We included two RT-qPCR kits QuantiNova and AgPath-ID One-Step RT-qPCR (ThermoFisher Scientific), two different platforms (CFX96 and LC480 I, (Roche)) and deviating primer (optimised and −33,3% (i.e. 0.7μM)) and probe (optimised and −50% (i.e. 0.2μM)) concentrations, erroneous reaction volumes (19μl or 21μl (+5μl of NA template)) and annealing temperatures of ±1°C (i.e. 63 and 65°C). In each combination the assay was tested using 10xLOD_95%_ of QS-1 in triplicates in a single run.

### Electrophoresis and sequencing

Electrophoretic analyses were performed on the TapeStation 4500 (D1000 Screen Tape and D1000 Reagents kits; all from Agilent Technologies) or by horizontal electrophoresis using a 2% (w/v) agarose gel (TAE buffer, and 10V/cm). Sequencing reactions were run using the Big Dye Terminator Cycle Sequencing Kit v3.1 and analysed on 3130 or 3500 Genetic Analysers (all from Life Technologies).

### Validation

The assay was validated through a combined effort of six veterinary or human diagnostic laboratories by using the following qPCR platform and chemistry combinations: i) CFX 96 (BioRad) and LC480 I (Roche)-QuantiNova Probe RT-PCR Kit (Qiagen); ii) LightCycler 480 II (Roche) and RotorGene Q (Qiagen)-OneStep RT-PCR (Qiagen) or QuantiNova Probe RT-PCR kits; and iii) Mx3000/5P thermocylers series (Agilent)-QuantiFast Probe RT-PCR+ROXvial Kit (Qiagen).

## Results

### Development of the SVIP-MPv2 assay

Re-assessment of the original SVIP-MP assay [24], which included a more recent (2018) analysis of 99,353 downloaded IAV MP-sequences [43] confirmed the high sequence conservation of the primers, guided the need for a new fluorescent probe design. Based on a highly conserved stretch at nucleotide positions 218-237, a novel FAM-labelled and MGB-modified probe SVIP-MP_P2-MGB, was selected (Table 1). This longer probe encompasses the original UPL104 probe motif with a K degenerate nucleotide at position 232, which improved assay inclusivity for contemporary human H3N2 strains (Supporting information 2, Figure S1).

**Figure 1.**
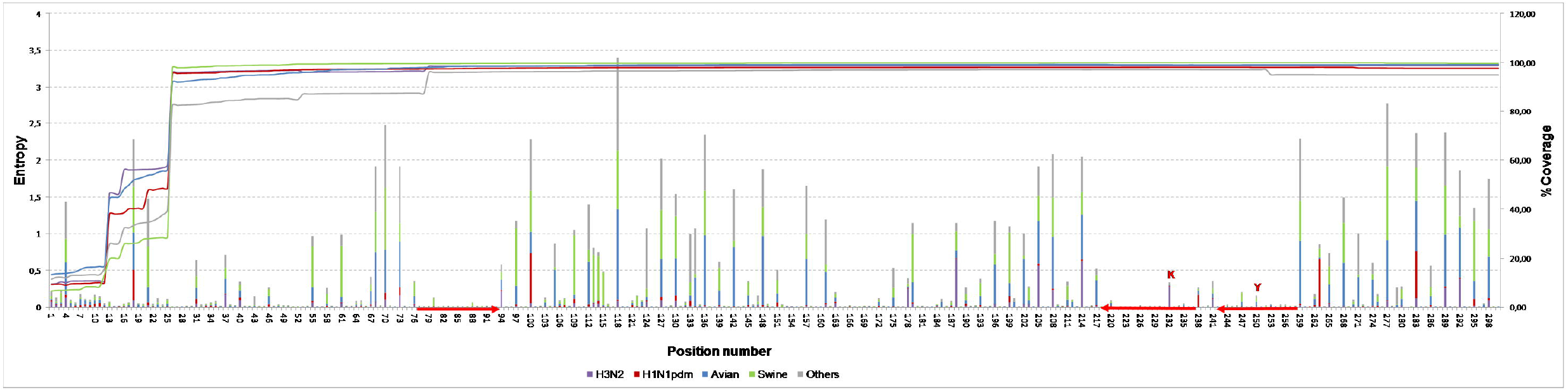
The entropy plot. The nucleotide variation of the IAV MP-segment within positions 1-300. The variability was calculated as entropy and was visualised separately for the five data pools: Human (H3N2 and H1N1pdm), avian, swine, and others, along the primary y-axis. The column heights are proportional to the amount of nucleotide variation/position observed. The line chart shows the coverage of the selected region in a given data pool, along the secondary y-axis. The primers and probe regions of the SVIP-MPv2 assay are highlighted in red. The PNNS of the primers and probe sequences are provided in Supporting information 2, Figure S1.

### Qualitative parameters of the SVIP-MPv2 assay

#### Efficiency

The calibration curve repeatedly showed strong linearity with a correlation coefficient R^2^≥0.996 and RT-qPCR efficiency ranged from 93-97%. Linear regression analysis inferred from raw fluorescent signals revealed mean efficiency of 1.94±0.06 (Supporting information 2, Figure S2).

#### Limit of detection

The sensitivity of the assay was expressed as LOD_95%_ i.e. the number of IAV genome equivalents detected with 95% confidence. The LOD_95%_ was estimated in two independent experiments by using dilution series of the QS-1 and QS-2 in a total genomic nucleic acid. The POD curve (Supporting information 2, Figure S3) demonstrated consistent detection of approximately ten IAV genomes per 25μl reaction.

#### Precision

The repeatability of the SVIP-MPv2 assay revealed SD of 0.33-0.40 and %RSD_r_ of 0.98-1.13% indicating absence of inherent random effects, even at highly diluted template quantities. The reproducibility showed SD of 1.04 and %RSD_R_ of 2.99% suggesting that the described SVIP-MPv2 protocol may be readily transferred to other laboratories.

#### Robustness

The six-parameter orthogonal combination matrix showed expected results with mean Cq of 35.76±1.18 and %RSD_R_ of 1.18% suggesting an acceptable standard of robustness. However, clear chemistry dependence was observed where the QuantiNova kit consistently showed lower Cq values in comparison with the AgPath kit. This was further confirmed by evaluating the SVIP-MPv2 assay with QuantiTect Probe RT-qPCR, OneStep RT-PCR (both from Qiagen) and AgPath RT-qPCR kits (data not shown). Finally, the robustness suggested independence from the qPCR thermal cycler model (data not shown).

### Validation of the SVIP-MPv2 assay

The SVIP-MPv2 assay validation included the specificity and inclusivity parameters.

#### Specificity

The specificity demonstrated the ability of the assay to selectively detect the desired template without any false positive signal. Two components of this parameter are recognized: i) Cross reaction with a host genomic nucleic acid background, or ii) with other microbial taxa present in a clinical specimen. The genomic interference was investigated on a total of 1358 IAV negative specimens of multiple matrices (Supporting information 1, AIV negative specimens and others). Potential cross reaction with other viral or bacterial genomes was investigated on a total of 409 specimens composed of 15 avian, five swine, three equine and 12 human viruses and 10 bacterial taxa. The human virus collection included also 35 SARS_CoV2 positive nasopharyngeal swabs. In addition, human origin field specimens positive for two non-IAV respiratory viruses in five different combinations were analysed. Finally, assay specificity was investigated on 12 different vaccines and 10 compound feed specimens for fattening poultry (Supporting information 1, others). In order to resolve weak positive FAM signals (high Cq values) and monitor the generation of nonspecific amplicons including primer dimers, the SVIP-MPv2 assay was run in MeltMan format [57]. The reactions with weak FAM curves (Cq>35) or melting peaks overlapping with the positive control were resolved by electrophoresis and if possible by amplicon sequencing. Positive FAM curves were always associated with the specific amplicon. We did not observe any non-specific, false positive FAM signal generation regardless of the host, virus or bacterial species and specimen matrix investigated including the taxonomically most closely related Influenza B (both Yamagata and Victoria lineages) and Influenza C Virus strains. This sufficiently proved the high specificity of the SVIP-MPv2 assay.

#### Inclusivity

This property is also known as the diagnostic sensitivity of the assay to detect as many subtypes, strains and phylogenetic lineages among all IAVs as possible. We distinguished two components of this parameter: Theoretical and experimental.

The theoretical inclusivity was calculated *in-silico* as the proportion of the analysed MP-sequences which has 100% identity to the primers and probe. To this end the sequence alignments were concatenated to include only the ROI and subsequently stratified. As seen, the concatenated human H3N2 data was decomposed to 39, the human H1N1pdm to 34, the avian to 112, the swine to 64 and the others to 40 ROI groups with highly asymmetrical frequency distributions (Table 2; Supporting information 2, Figure S4). Only the ROI groups situated above the empirical threshold of 0.5% were used for the theoretical inclusivity estimation. As a result, 100% of human (both H3N2 and H1N1pdm), 98.2% of avian, 94.6% of swine and 100% of other IAV strains exhibited perfect sequence identity to the primers and probe sequences. Overall, this resulted in a 99% detection rate at the defined threshold level. The concatenated alignments are available as a Supporting dataset, enabling all of the primers or probe sequence variants to be identified by the SequenceTracer web tool [43].

The experimental inclusivity demonstrated the real ability of the assay to universally detect various IAV strains, subtypes and pathotypes in a spectrum of specimen matrices collected from different host species. The inclusivity of the SVIP-MPv2 assay was evaluated on a total of 612 IAV positive specimens of 50 subtype combinations. The list of all IAV strains, subtypes and pathotypes as well as collection dates and specimen matrices are listed in Supporting information 1. All of the IAVs were successfully detected at the participating laboratories, which was in full agreement with the predicted expectations and proved the high inclusivity of the SVIP-MPv2 assay.

Experimental validation at APHA (UK) of the SVIP-MPv2 assay included parallel testing of a wide range of AIV isolates (n=77) by the earlier SVIP-MPv1 [24] and the M-gene RT-qPCR [16] assays (Supporting information 3). Similar parallel testing of SwIV isolates (n=54) and seven porcine respiratory tissues involved substitution of the M-gene “perfect match” RT-qPCR [36] for the Spackman assay [16]. The APHA (UK) validation also included the testing of 79 swabs obtained from AIV outbreaks and experimentally-infected birds, assessed by both the SVIP-MPv1 and v2 RT-qPCRs. Overall, the SVIP-MPv2 assay demonstrated similar Cq values to those registered by the other M-gene RT-qPCRs (Supporting information 3).

### Identification of potentially critical IAV variants

The inclusivity assessment revealed three potentially critical ROI variants in the avian (Figure 2, groups 3-5) and three in the swine MP-sequence data (Figure 2, groups 2, 3 and 5). Especially, the avian-groups_3-5 and swine-groups_3 and 5 are of concern due to potential interference with probe annealing or 3’-priming. For a detailed investigation the Fasta headers were extracted from the corresponding MP-sequence groups and the spacio-temporal characteristics were further investigated. Temporal distribution chart (Figure 3) revealed three sharp peaks corresponding with avian-group_4-reverse, swine-group_2-forward and swine-group_5-reverse primers. The avian-group_4-reverse consists mainly of A/mallard/northern_pintail/(H3N8, H4N6, H10N7) strains from Alaska from a single sampling period in 2009. The swine-group_2-forward and swine-group_3-reverse were both among the US swine strains (all subtypes, i. e. H1N1, H1N2, H3N2) obtained between 2012-2017 and 2011-2016 respectively. The remaining groups were distributed at a low frequency since 2000-2003 with a slight peak of the avian-group_3-probe in 2015 (mainly A/duck/chicken/goose/Hunan/2015(H5N6)-like strains) and the swine-group_5 in 2014-2015 (predominantly A/swine/US/2014 and 2015 strains of all three subtypes). However, all of these variants occurred within the background population of the prevailing perfect match strains which dominated at the same time in a given sampling locality (Figure 3) as determined from the submitted sequence data.

**Figure 2.**
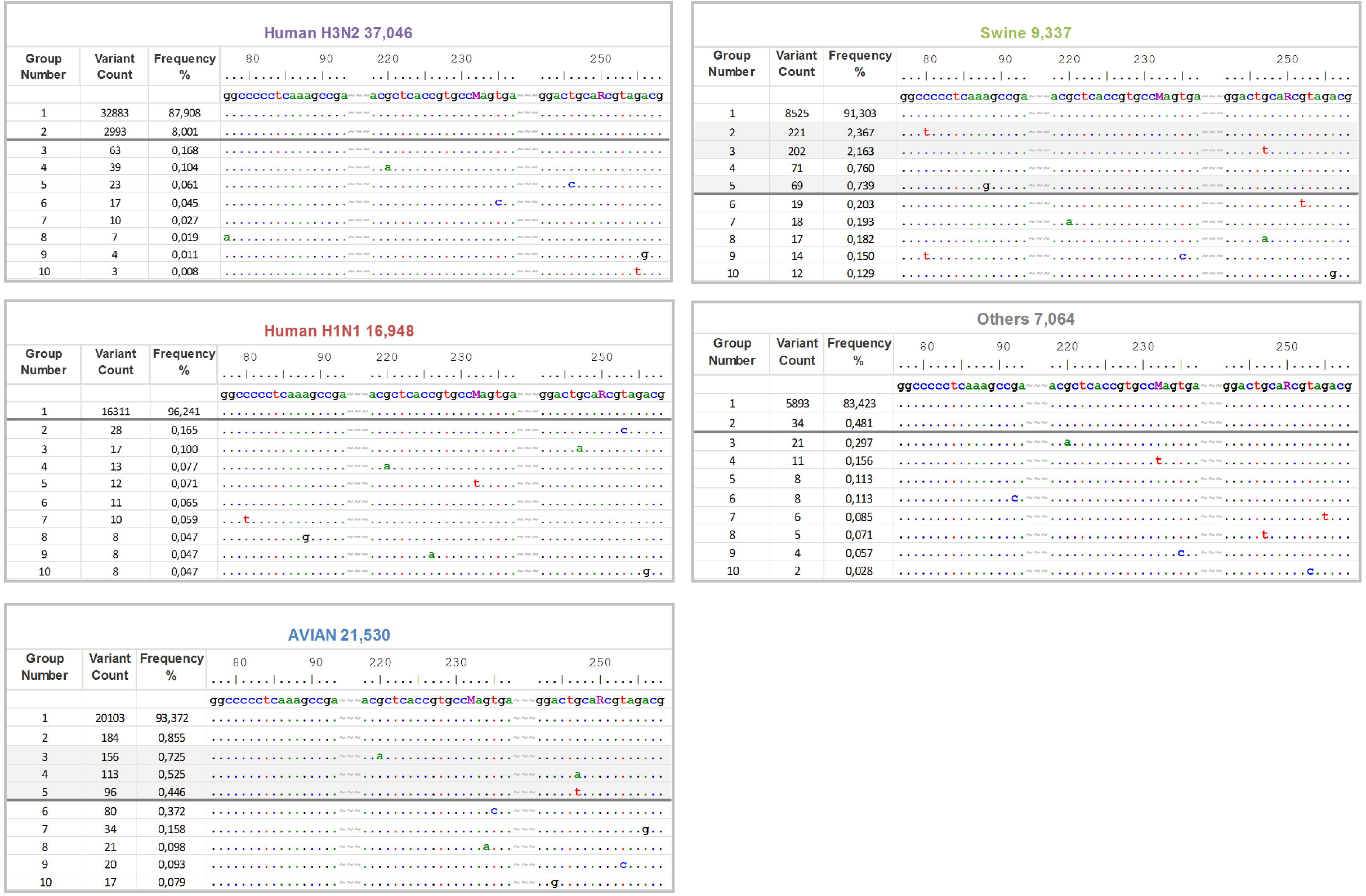
Alignment stratification of the data pools. The concatenated (forward-probe-reverse) alignments of the human H3N2 and H1N1pdm, avian, swine, and other IAVs prepared by the SequenceTracer web tool. The alignment shows the first ten M-sequence groups in a descending order aligned to the SVIP-MPv2 primers and probe sequences (5’→3’). Note that the probe and the reverse primer sequences are shown in the forward (positive stranded) orientations. The alignment was drawn in the Graphic View mode of the BioEdit program. For clarity, the primers and probe sequences were separated by tildes and numbered according to their MP-segment positions. The dots indicate nucleotide identity and the horizontal grey bars designate the threshold (≥0.5%). The shaded groups designate potentially critical MP-segment variants. For full alignment stratifications please refer to Supporting information 2, Figure S4.

**Figure 3.**
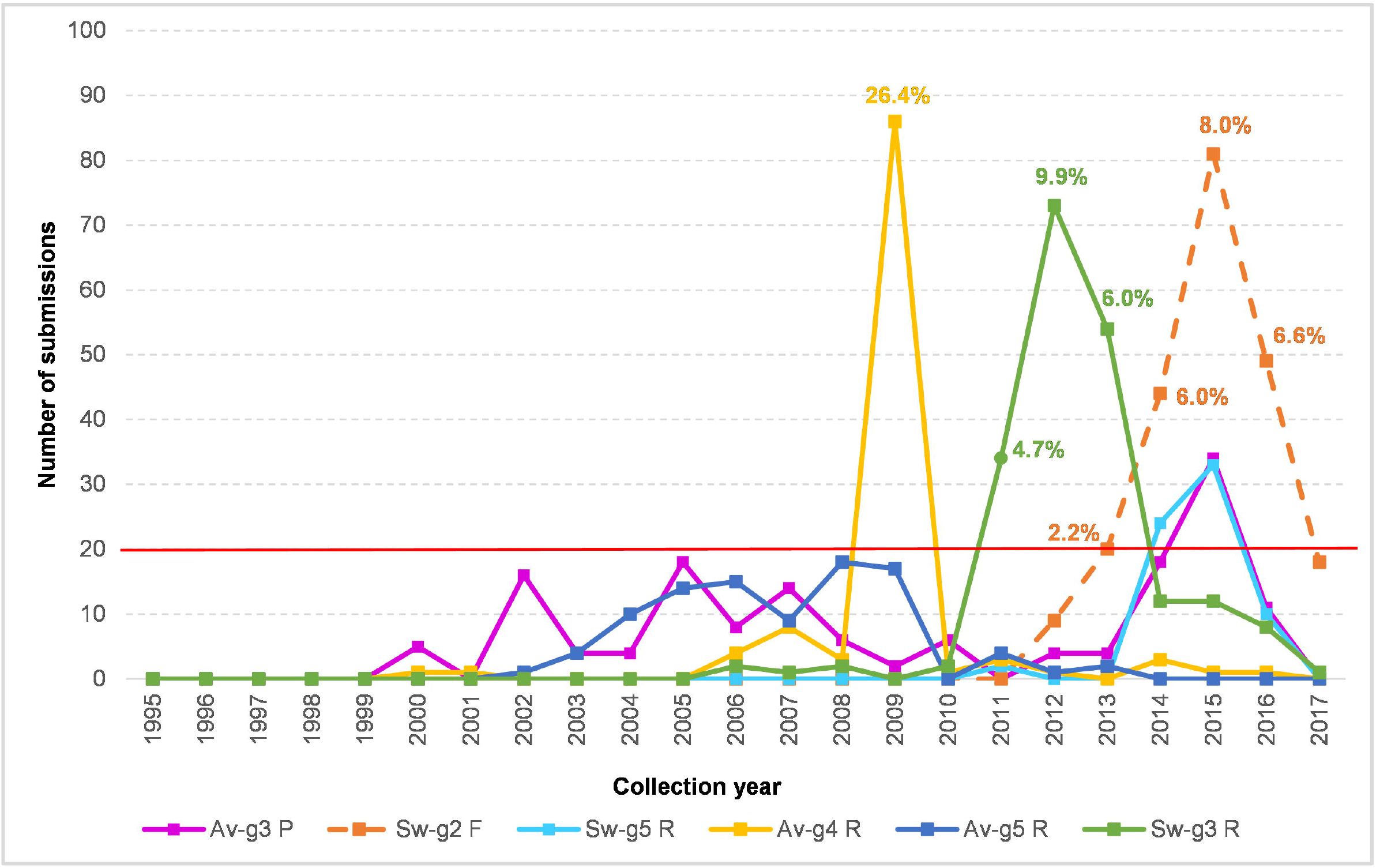
Temporal distribution chart of the critical ROI variants. The Fasta headers were extracted from the avian-groups_3-5 and swine-groups_3 and 5 (Figure 2; shaded) by the FaBox web tool and the submission counts per year were visualised as a line chart. The proportion of the particular group in the submissions available from the same year and geographic area were shown at key nodes as percentages.

## Discussion

The study successfully exploited thorough bioinformatics analyses of the IAV MP-segment to modify an earlier RT-qPCR design by simply altering the probe sequence. Our aim was to provide a new version of the assay, SVIP-MPv2, which compensated the deficiency observed for the original version [43] and now meets the requirement for multi-species origin IAV detection in accord with the One Health initiative. In addition, the new version is independent of the proprietary commercial UPL104 probe employed in the first version, where the position, number and character of the LNA modified nucleotides have not been disclosed. There have also recently emerged difficulties in the future procurement of the UPL104 probe, so without open knowledge of its full sequence details there is presently little or no possibility of an alternative supplier.

Despite the challenges posed by the aims of this study, the SVIP-MPv2 assay was effectively optimised with a LOD_95%_ of approximately ten IAV genome templates per reaction with high repeatability, reproducibility and robustness. The assay is compliant with the MIQUE guidelines [54] and the parameters fulfil the crucial diagnostic RT-qPCR criteria and validation guidelines [55]. Although these guidelines were intended for detection of genetically modified organisms, several approaches such as LOD and robustness were also adopted in our study to underline the importance of method performance and criteria for acceptance. However, the main advantage of the SVIP-MPv2 assay is its high inclusivity in detecting a range of diverse-origin IAVs. The *in-silico* analysis predicted 99% IAV detection rate in the analysed collection of MP sequences (n= 99,353), which included all known IAV subtypes from various host species and geographical origins obtained since 1902 [43]. The IAV-like subtypes H17N10 and H18N11, described exclusively in bats [59, 60], were excluded from the analysis because they do not pose any threats to human or livestock health and are also genetically highly distinct from other IAV subtypes.

The theoretical inclusivity predictions were confirmed experimentally by six veterinary or human public health diagnostic laboratories which independently tested diverse panels of human, avian, swine and equine IAV strains obtained in a range of specimen matrices. Testing of IAV negative specimens demonstrated the high specificity of the SVIP-MPv2 assay, and included other avian viral pathogens of note. Validation included testing AIVs of diverse origins which included all 16 HA and nine NA subtypes. The AIVs included isolates and clinical specimens obtained from the most recent incursion of the important GsGd lineage into Europe, namely the H5N8 HPAIV, during early 2020 [61]. Seventy-seven AIV isolates were tested by the SVIP-MPv2 assay alongside the earlier validated SVIP-MPv1 [24] and Spackman et al. [16] M-gene RT-qPCRs with an overall close concordance evident among the recorded Cq values, which was also observed between the first two of these assays when applied to the testing of 79 swabs obtained from AIV outbreaks or from experimentally-infected birds (Suppl information 3). The choice of well-characterised IAV positive and negative specimens, both laboratory isolates and those of clinical origin, reflected the OIE guidelines for validation which underlined the need to include such reference samples [35] (OIE, 2018a). The SVIP-MPv2 test also detected all IAV strains in the proficiency test organised by the European Union Reference Laboratory for AIV (Instituto Zooprofilacttico delle Venezie, Italy) in 2019 (data not shown).

Replacing the probe clearly improved the *in silico* inclusivity of the SVIP-MP assay for the human H3N2 IAVs to 100% relative to the original 91.7% [43]. Sensitive H3N2 detection is important not only for public health, but also because this human subtype can transmit into swine leading to the establishment of new reassorted lineages [11,62, 63]. Importantly, the assay sensitively detected the SwIV genotypes which contain the internal gene cassette of H1N1pdm origin which have become increasingly more prevalent during the past decade, regardless of the subtype determined by the glycoprotein (external) genes [11, 64]. As in the case of validation of the SVIP-MPv2 assay with well-characterised AIV isolates and clinical specimens, parallel testing of SwIVs was carried-out alongside the earlier validated M-gene RT-qPCRs, namely the SVIP-MPv1 [24] and ‘perfect-match’ assays [36]. Again, an overall close concordance was observed among the recorded Cq values for the SwIV isolates (n=54) and clinical specimens (n=10).

Returning to the importance of the new SVIP-MPv2 probe design based on bioinformatics analysis the new probe retained unchanged inclusivity for the various data pools, i.e. 100% for human H1N1pdm, 98.2% for avian, 94.6% for swine and 100% for others assessed at the 0.5% threshold level. Consequently, the overall inclusivity rate increased to 99.0% from the original 95.8%. On the other hand, the slight decrease in the inclusivity for the swine IAVs (94.6%) was mainly due to the variations in the reverse primer binding positions of the US SwIVs, albeit at a low background frequency. Therefore, despite being extremely conserved, the reverse primer represents potentially vulnerable component of the assay among the US swine IAV MP-sequences. Unfortunately, the SwIVs included in the experimental validation did not include the corresponding MP-segment variants, therefore the SVIP-MPv2’s ability to detect these variants was not investigated. This small concern warrants the ongoing assessment of primers and probe sequences *in silico* in order to monitor for any sustained mutational trends in this region. Nevertheless, the assay’s high inclusivity for various species combined with high sensitivity predestines the SVIP-MPv2 as a universal IAV screening tool, not only for the One Health species, but also for minor or occasional mammalian hosts which merit investigation as a risk for zoonotic infection [65, 66]. The careful design of the SVIP-MPv2 assay has taken into account the long term evolutionary trends which continue within conserved genes such as MP1, so reducing the need for future modifications or adjustments which were required for the assay of Spackman et al., [16] to provide detection of pandemic H1N1 viruses in swine [36] or H5 HPAI viruses in wild and domestic birds [23, 67, 68]. Similarly, it can also reduce the number of different generic RT-qPCR techniques recommended for IAV surveillance in the human population [37].

Large-scale multiple sequence alignments (LS-MSA) represent an invaluable information source for a universal diagnostic PCR design or reassessment [69–71]. A comprehensive analysis of sequence variability based on LS-MSAs was also critical to the presented study. Nevertheless, sequence databases may be vulnerable to errors and biases which are important to consider. Although errors such as single nucleotide insertions, redundant stretches of primers or cloning vectors, sequence submissions in incorrect directions etc. can be recognized relatively easily [51,72] and correctable during alignment, but others may be more difficult to identify [73]. In addition, it is important to consider the compositional bias which addresses significant under- or over-representation of certain sequence variants. For instance, 55% of the avian primer-probe-primer groups included only a single MP-sequence variant. How many of these may be a sequencing error, thereby representing a degree of misleading data? Conversely, how many of these are true variants, reflecting hidden evolutional plasticity of the MP-segment and potential for future emergence as a new MP variant? While some compositional bias of the data is unavoidable the variants with the lowest frequency were discarded by setting a threshold of 0.5%.

Generally, a RT-qPCR design for universal IAV detection represents a difficult challenge. The predefined lengths and sequence motifs of the conserved regions greatly limit the freedom for oligonucleotide selection [69]. Other than perfect fit to the conserved regions, the selected primer and probe candidates may not exhibit ideal thermodynamic parameters such as sufficient lengths, Tm values or GC content. In addition, oligonucleotides may be vulnerable to a degree of homo- or hetero-dimerization, or hairpin formation. To overcome these undesired properties, the SVIP-MPv2 assay had to be rigorously optimised, which included adjustment of the annealing temperature to 64°C together with increased primers and probe concentrations to achieve optimal reaction parameters.

The history of IAV outbreaks suggests that novel epizootic or pandemic strains can emerge suddenly from a previously unrecognized virus population. If the new IAV variant were to include undesirable primer or probe mutations, its emergence could compromise diagnostic issues especially during the early stages of the outbreak when prompt responses based on accurate epidemiological or field information are critical for the implementation of appropriate control measures. Therefore, sensitive and specific detection of the broadest range of different IAV strains and subtypes, irrespective of the host source, is of utmost importance. This study has provided a holistic approach to the development and evaluation of a RT-qPCR assay for universal IAV detection. The primers and probe, selected on the evolutionarily ultra-conserved regions in the MP-segment, were a crucial consideration to ensure the assay’s broad inclusivity for detection of the diverse IAV range. In conclusion, the modified MP-segment RT-qPCR should be used as a frontline screening tool to detect diverse IAV strains, particularly when considering the One Health perspective.

## Supporting information

Summary of all host species, non-IAV and bacterial taxa as well as IA virus subtypes, pathotypes, collection dates and specimen matrices included in t

Qualitative reaction parameters of the SVIP-MPv2 RT-qPCR assay.

Validation of the SVIP-MPv2 at APHA-Weybridge (UK) using avian, swine and human IAV isolates and clinical specimens from AIV- and SwIV-infected birds

The concatenated primer-probe-primer sequences for avian, swine, equine, human and other data.

## Acknowledgements

This work was dedicated to Dr. Martina Havlíčková. head of the National Reference Laboratory for Influenza, National Institute of Public Health, Czech Republic, our friend and colleague, who passed away in December 2019. We thank all contributors to the EpiFlu (Global Initiative on Sharing All Influenza Data) and IVD (National Centre for Biotechnology Informations) databases. Further, Drs Kristien Van Reeth (Univerity of Gent, Gent, Belgium) and Gustavo del Real (Instituto Nacional de Investigación y Tecnología Agraria (INIA) Madrid, Spain) are gratefully acknowledged for their permission to include testing of SwIV isolates from their respective countries.

## Author contributions

AN: conceptualization, data curation, formal analysis, investigation, methodology, project administration, software, supervision, validation, writing-original draft preparation

LČ: investigation, methodology, validation, probe selection

KK: investigation, methodology, validation

ZD: investigation, methodology, resources, validation

SST: data curation, investigation, resources, validation, visualization, writing-review and editing

MJS: funding acquisition, project administration, resources, supervision, validation, visualization, writing-review and editing

ÁD: formal analysis, investigation, methodology, supervision, validation

TV: investigation, methodology, validation

MM: investigation, methodology, validation

HJ: investigation, methodology, validation

IHB: funding acquisition, resources, writing-review and editing

All authors approved the final manuscript.

## Funding

State Veterinary Institute Prague, Czech Republic and Veterinary Institute Zvolen, Slovak Republic: The work was performed in connection with the activities of the National Reference Laboratory for Avian Influenza Viruses and was not funded by additional financial sources.

Animal and Plant Health Agency, United Kingdom: The work was funded by the Department of Environment, Food and Rural Affairs (Defra, UK) and the Devolved Administrations of Scotland and Wales through (i) the SE2211 Programme of Work “FLUFUTURES - Reducing Influenza A Virus (avian and mammalian) hazards in the UK”; plus two Defra Contract C programmes, namely (ii) SV3400 National Reference Laboratory for Statutory and Exotic Virus Diseases of Avian Species and Relevant Hosts, and (iii) SV 3006 for the International Reference Laboratory for Avian Influenza and Swine Influenza.

Danam.Vet.Molbiol, Budapest, Hungary; SCG Diagnosztika Ltd., Délegyháza, Hungary; Prophyl Animal Health Ltd., Mohács, Hungary: The companies own budgets were used. No funding by any additional financial sources.

National Institute of Public Health, Czech Republic: The work was supported by Ministry of Health, Czech Republic - conceptual development of research organization (,,The National Institute of Public Health – NIPH, 75010330“).

## Conflicts of interest

The authors declare that they have no conflicts of interest.

## Supporting information captions

**Supporting information 1:** Summary of all host species, non-IAV and bacterial taxa as well as IA virus subtypes, pathotypes, collection dates and specimen matrices included in the study.

**Supporting information 2:** Qualitative reaction parameters of the SVIP-MPv2 RT-qPCR assay.

**Supporting information 3:** Validation of the SVIP-MPv2 at APHA-Weybridge (UK) using avian, swine and human IAV isolates and clinical specimens from AIV- and SwIV-infected birds and pigs respectively.

**Supporting dataset:** The concatenated primer-probe-primer sequences for avian, swine, equine, human and other data.

## Notes

### Competing Interest Statement

The authors have declared no competing interest.

