## Supplementary material for "A universal RT-qPCR assay for “One Health” detection of influenza A viruses": Qualitative reaction parameters of the SVIP-MPv2 RT-qPCR assay.

**Supporting Information 2.** Qualitative reaction parameters of the SVIP-MPv2 RT-qPCR assay.

**Figure S1. PNNS for the primers and probe sequences in the five data pools.**

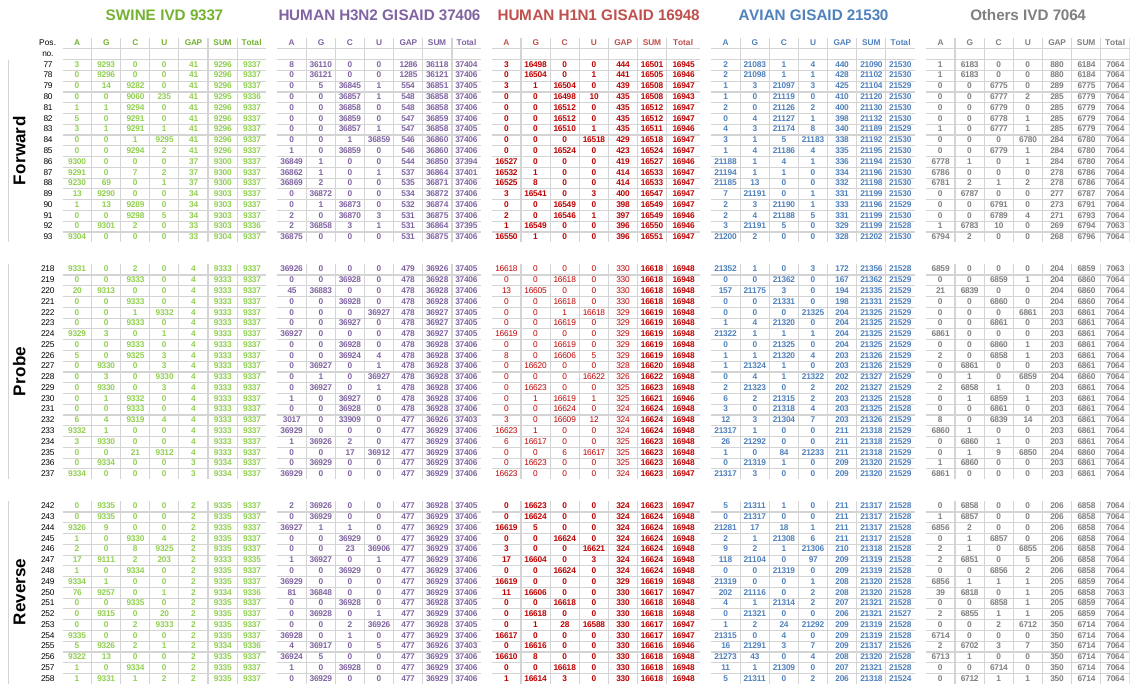

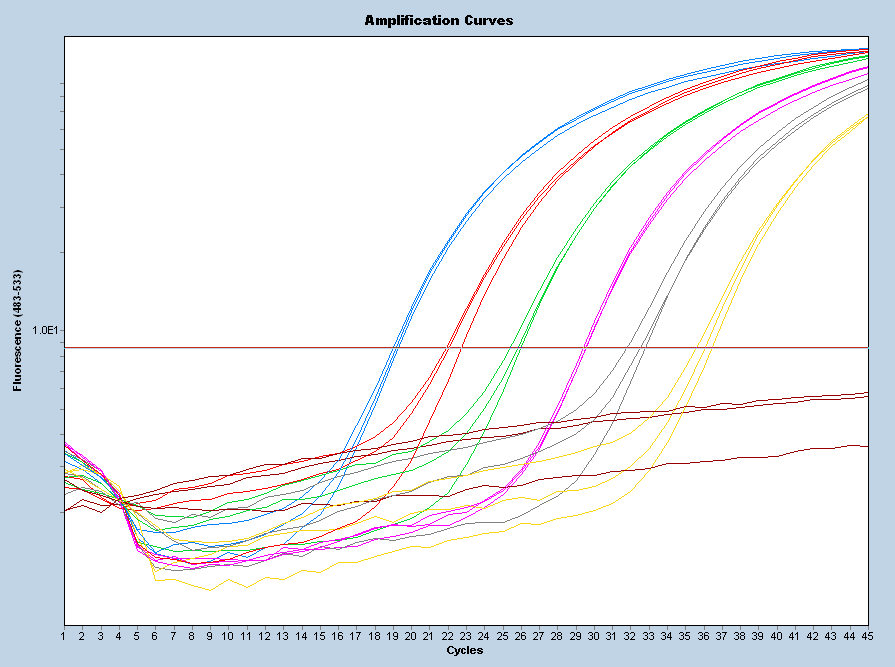

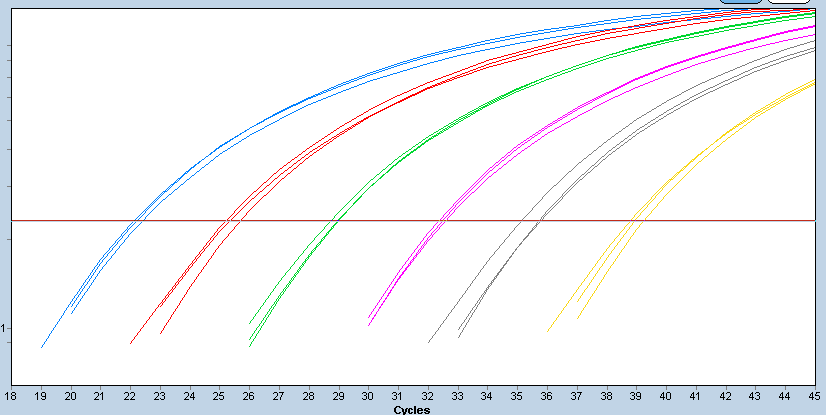

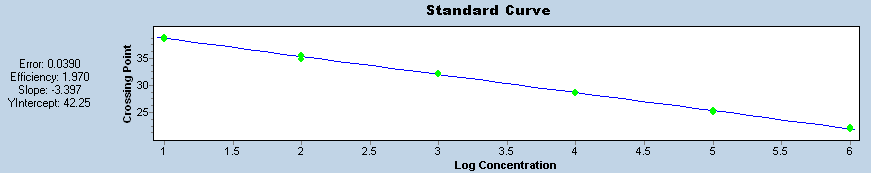

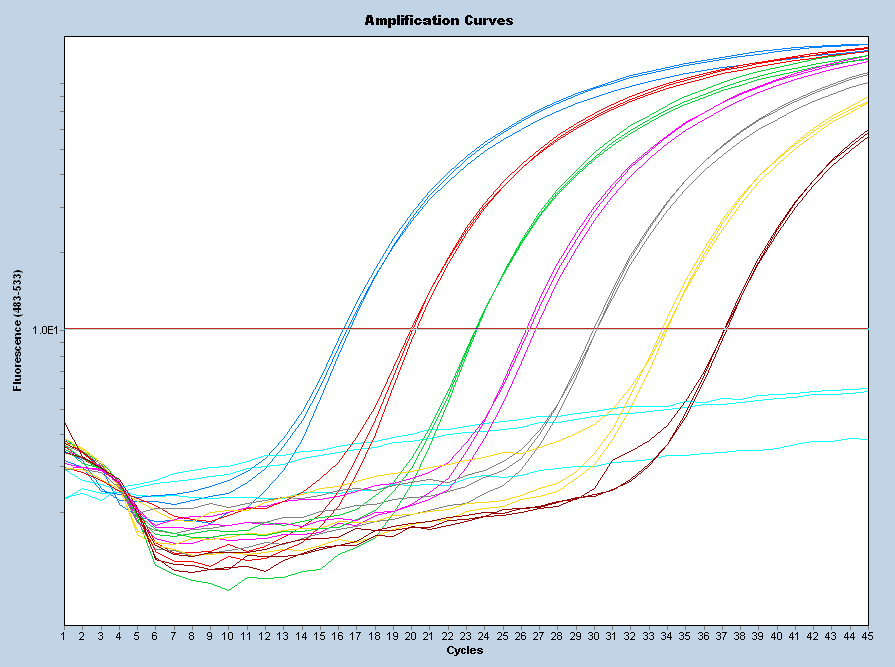

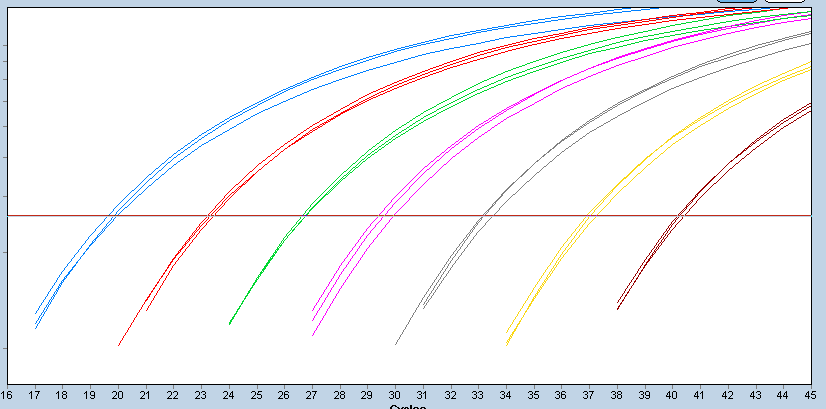

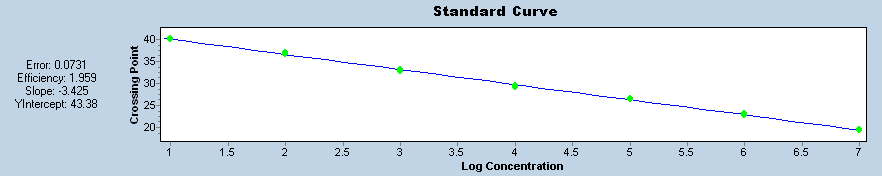

A/mallard/Czech Republic/13579-84K/2010(H4N6) [JF789621](http://www.ncbi.nlm.nih.gov/entrez/viewer.fcgi?val=JF789621) 1e6-1e1 copies/μl

A/goose/Czech Republic/1848-T14/2009(H7N9) [HQ244418](http://www.ncbi.nlm.nih.gov/entrez/viewer.fcgi?val=HQ244418) 1e7-1e1 copies/μl

**Figure S2. Efficiency estimation of the SVIP-MP2 assay.** Calibration and standard curves were constructed by 10-log dilution of the QS-1 and QS-2 in a concentration from 1e6 to 1e2 (QS1) or from 1e7 to 1e2 (QS-2) copies/µl in triplicates. The dilutions were prepared in a total background nucleic of swine origin (at least 7ng/µl) and the results are shown for the LC480 instrument.

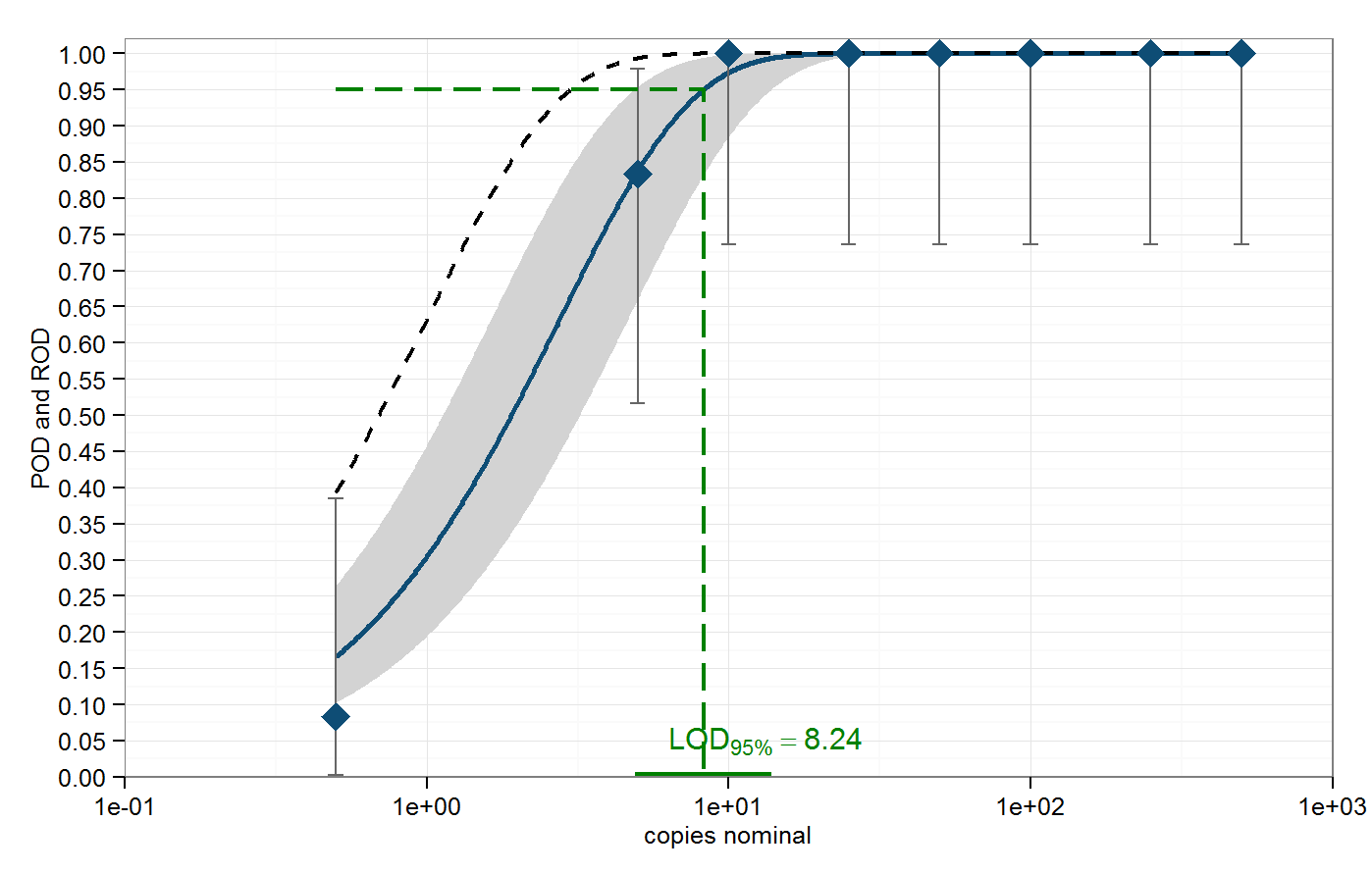

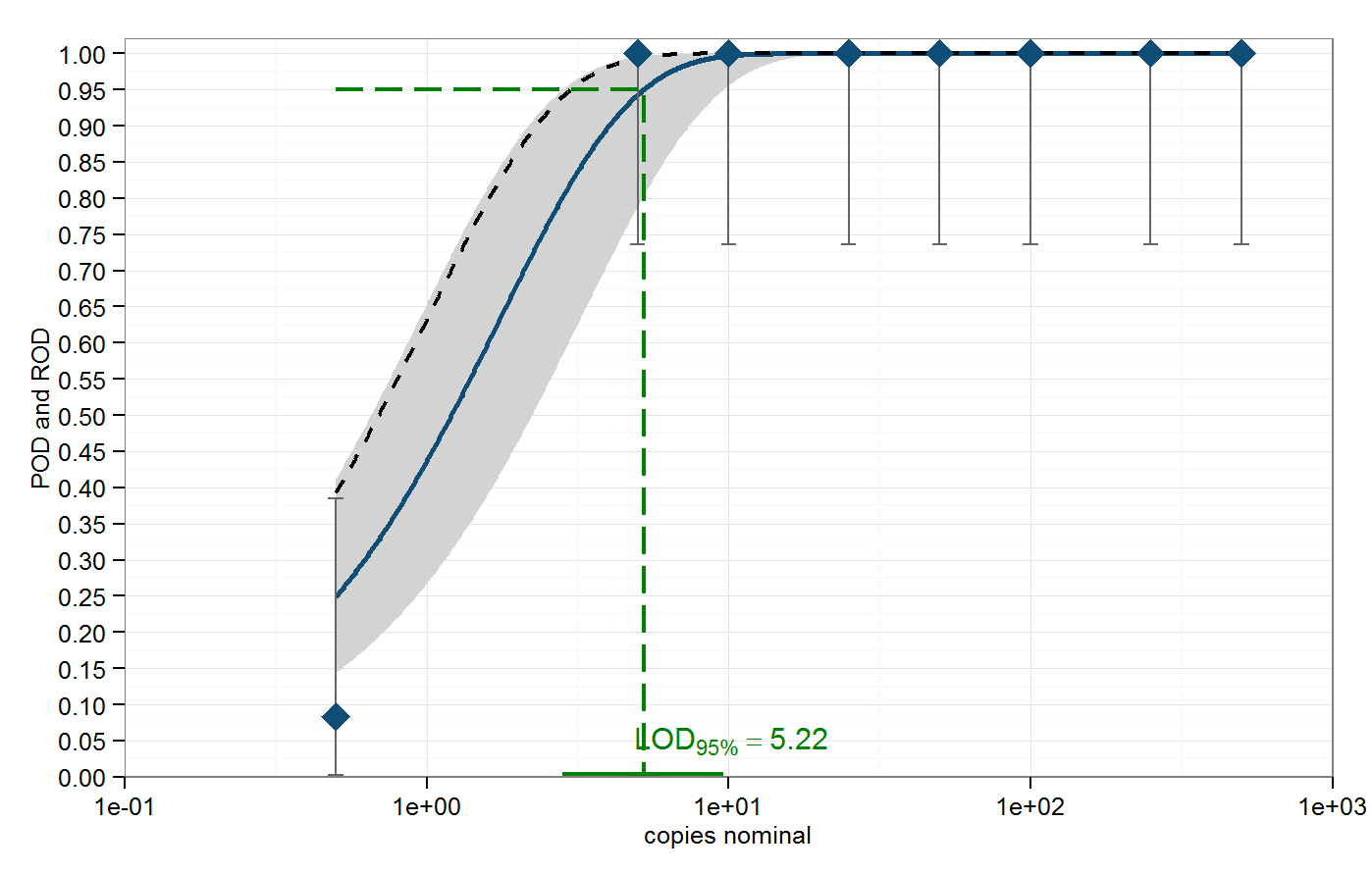

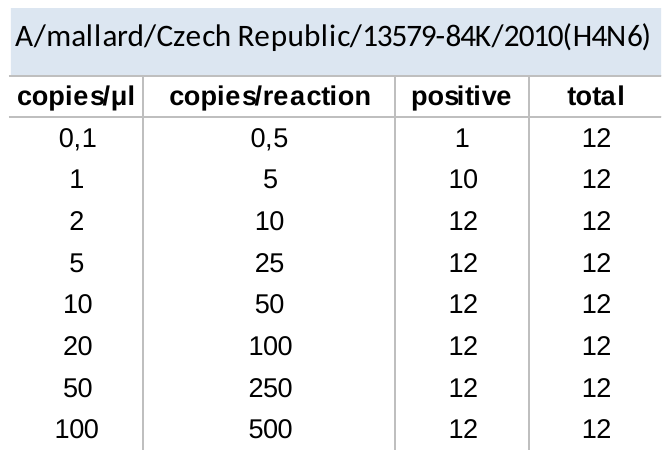

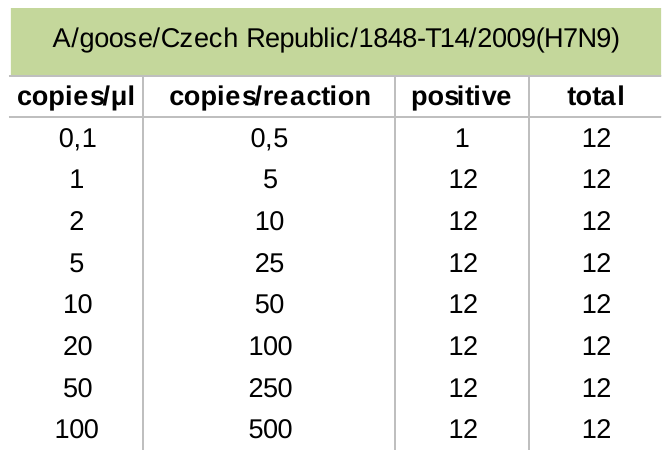

**Figure S3. Limit of detection.** The probability of detection (POD) curves and LOD_95%_ for (A) QS1 (strain A/mallard/Czech Republic/13579-84K/2010(H4N6); GenBank accession number [JF789621](http://www.ncbi.nlm.nih.gov/entrez/viewer.fcgi?val=JF789621)) and (B) QS2 (strain A/goose/Czech Republic/1848-T14/2009(H7N9; GenBank accession number [HQ244418](http://www.ncbi.nlm.nih.gov/entrez/viewer.fcgi?val=HQ244418)). The blue curves denote the mean POD curve along with the corresponding 95 % confidence range highlighted in grey. The POD under ideal conditions is displayed as the black dashed curves. The LOD_95%_ for the QS1 was established as 8.243 copies/reaction, i.e. ~2 copies/µl of template with a 95 % confidence interval of [4.897, 13.913] and for QS2 as 5.220 copies/reaction, i.e. ~1 copy/µl of template with a 95 % confidence interval of [2.817, 9.644]. The tables (C, D) show the positive rate obtained per 12 replicates (2*6 replicates under repeatability conditions).

**C**

**A**

**D**

**B**

**Table S1.** The orthogonal combination matrix scheme (A) along the factors and their values (B) considered for robustness estimation**.**

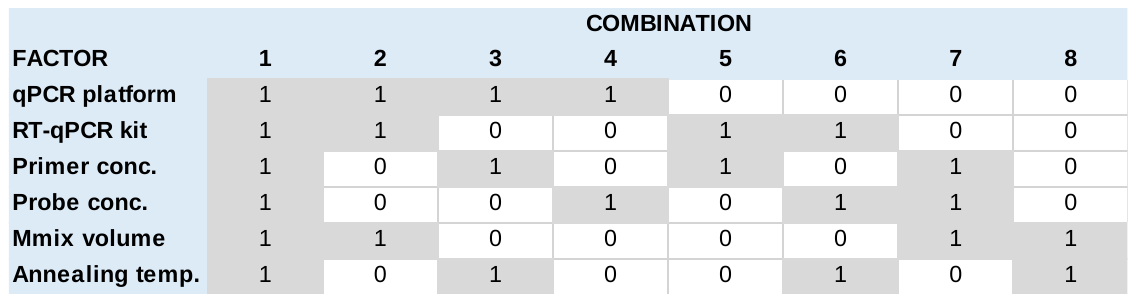

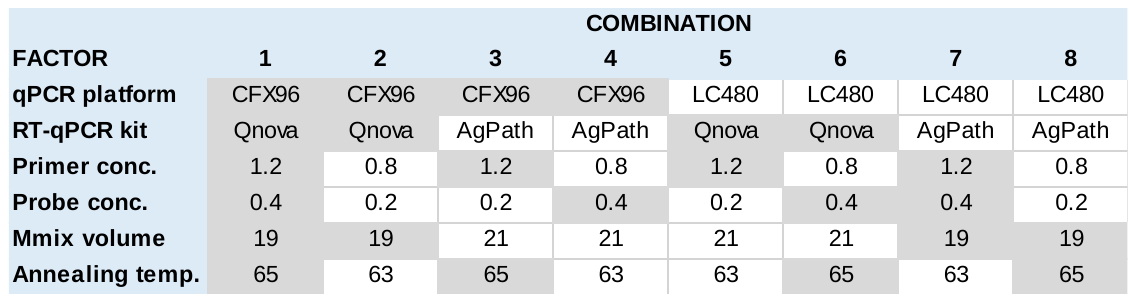

**A**

**B**

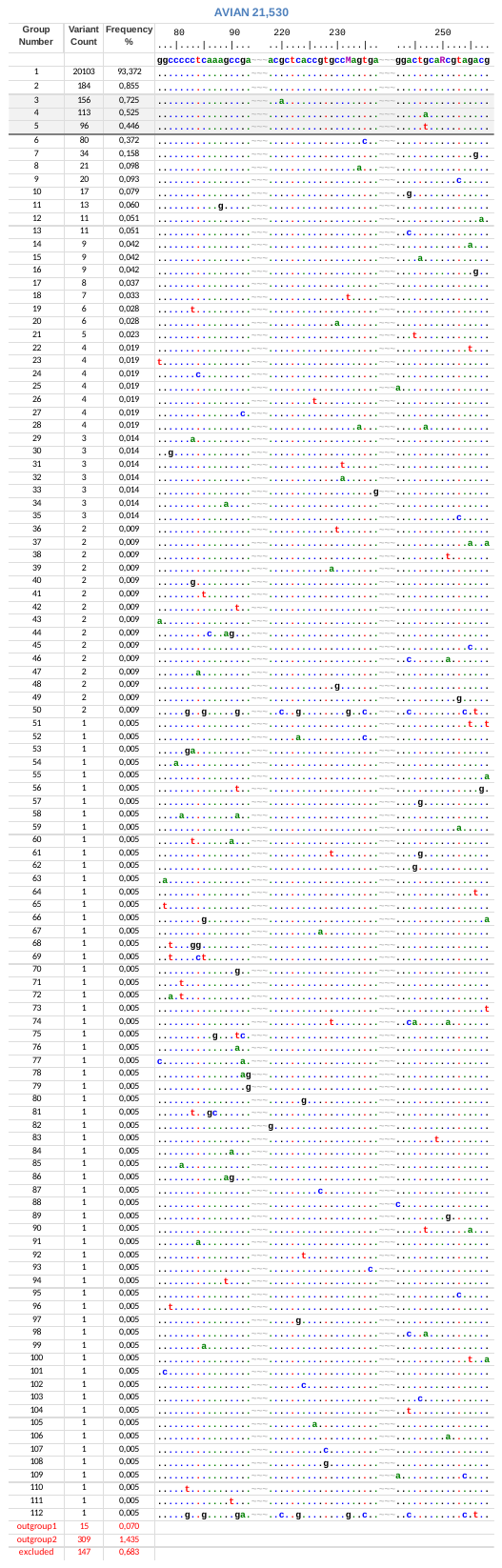

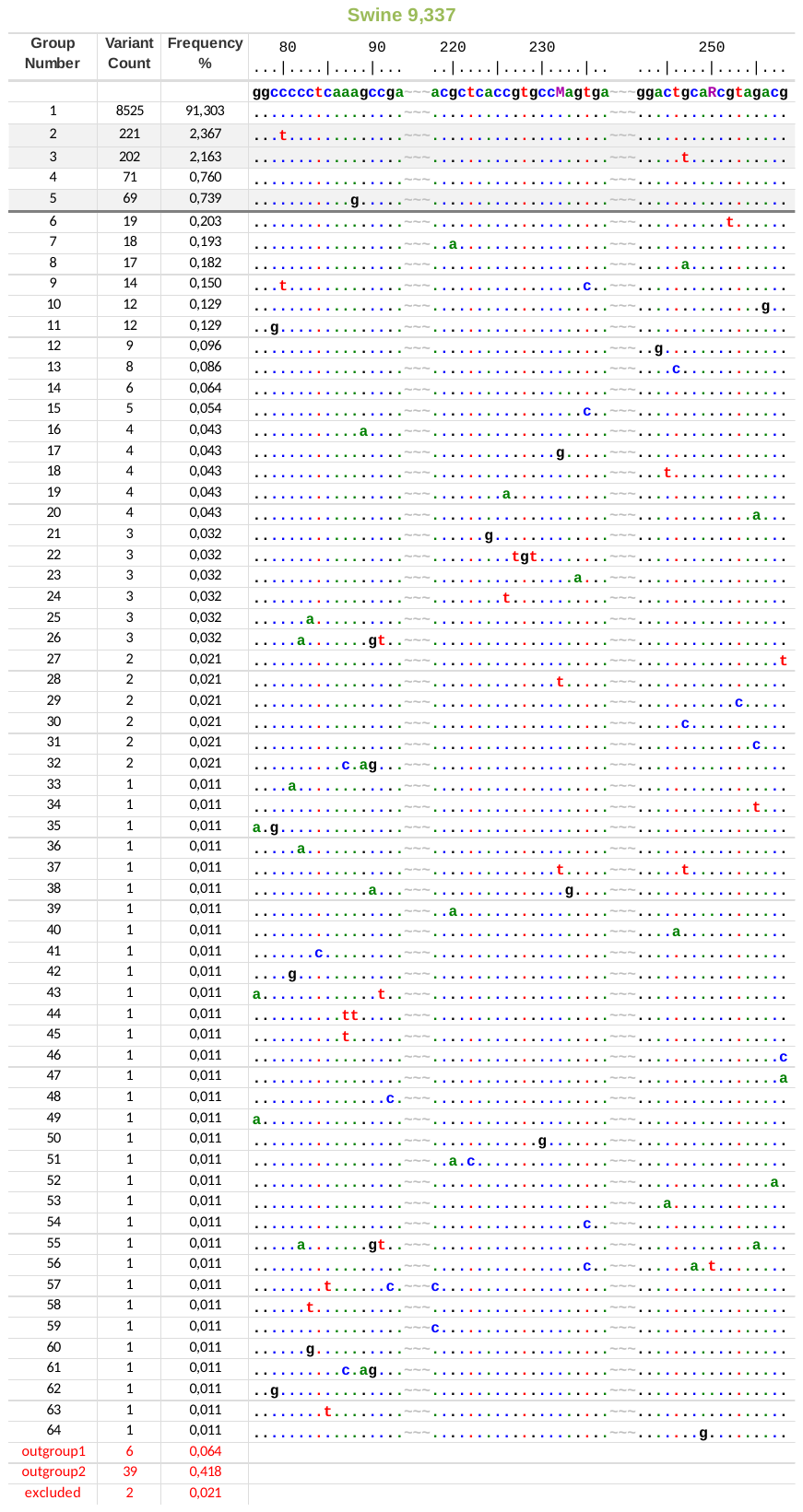

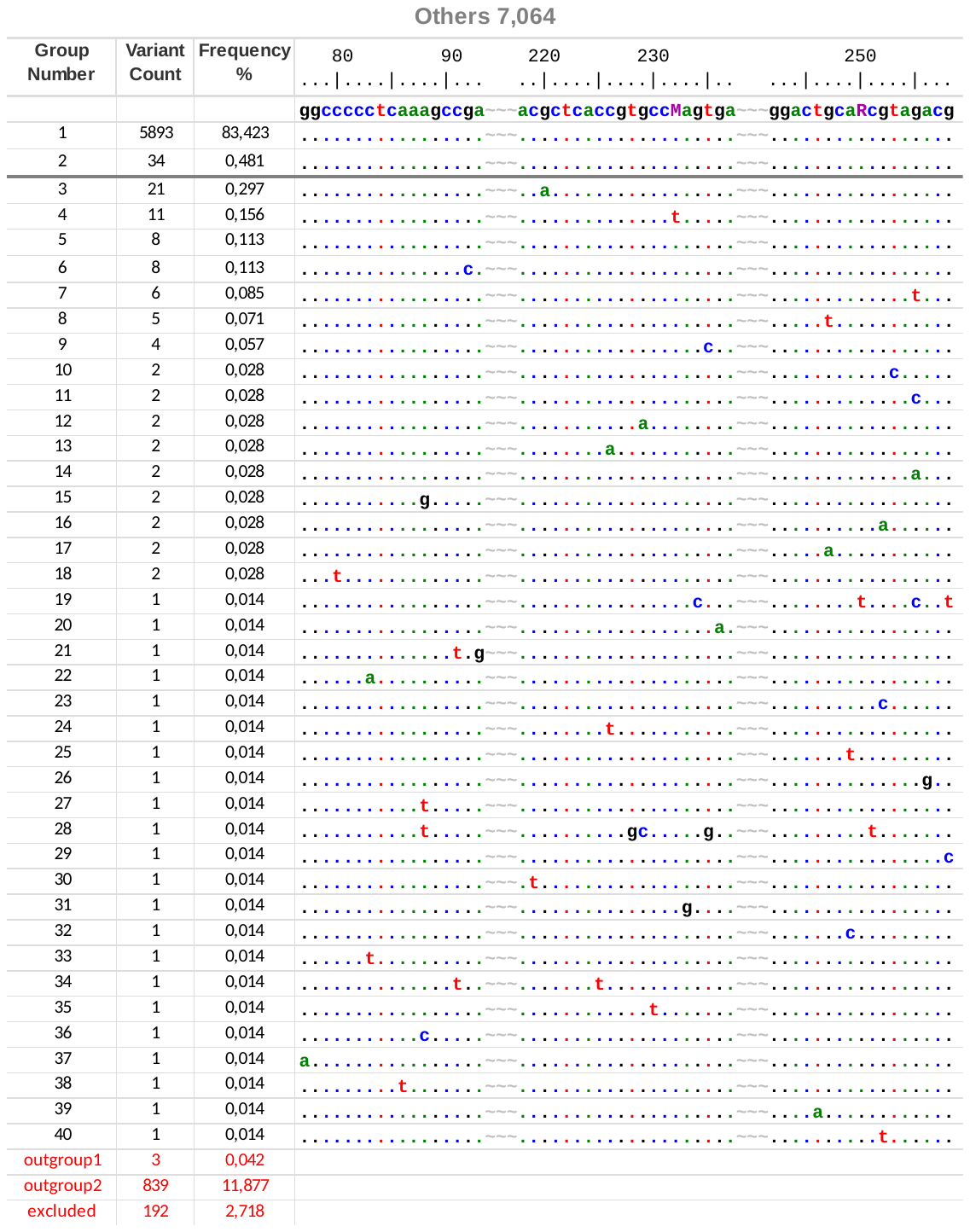

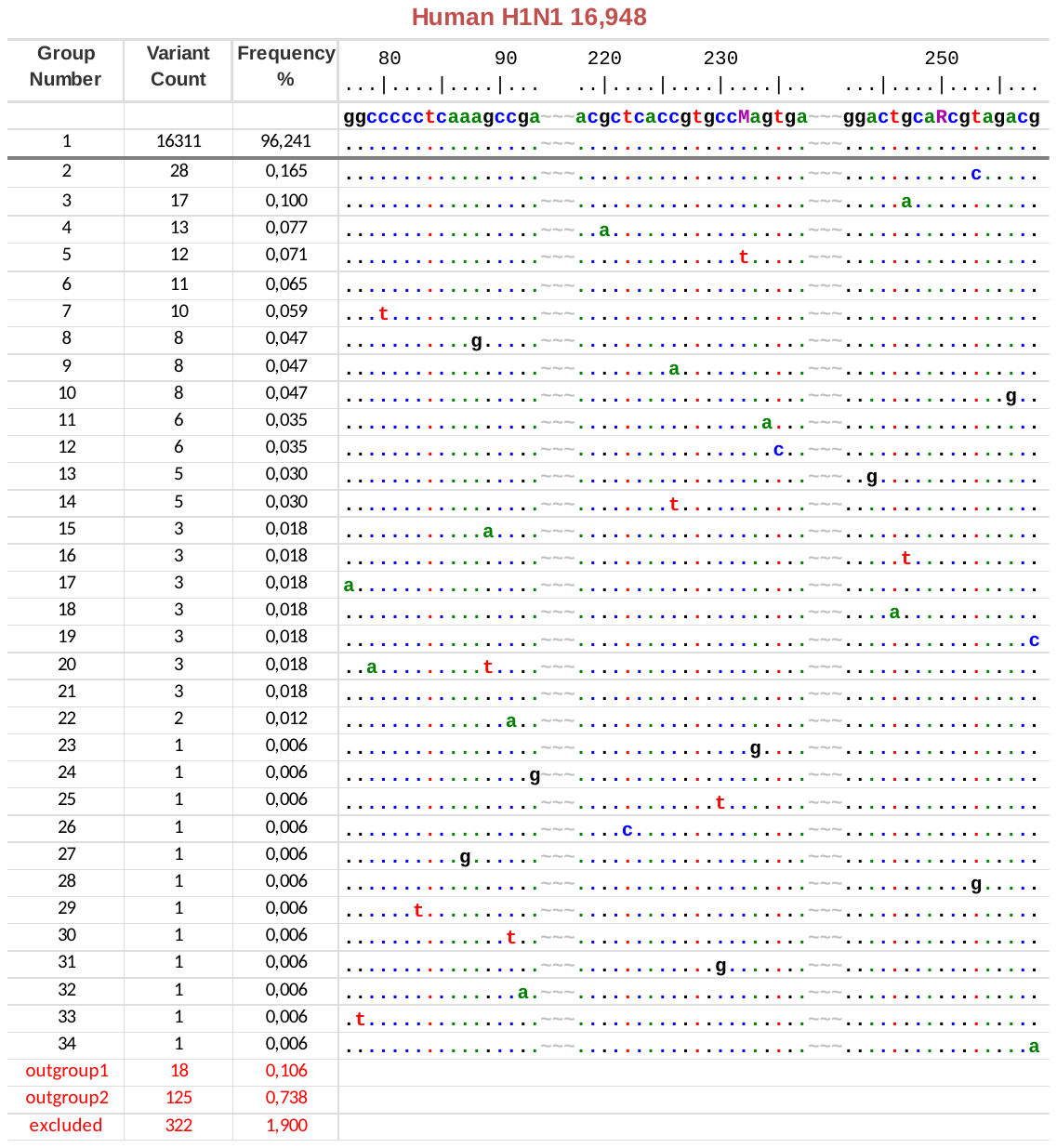

**Figure S4. Alignment stratification.** The concatenated (forward-probe-reverse) alignments of the human H3N2, human H1N1pdm, avian, swine, and others data prepared by the SequenceTracer web tool. The figure shows all the M-sequence groups in a descending order along with a representative sequences aligned to the SVIP-MPv2 primers and probe sequences (5’→3’). Note that the probe and reverse primer sequences are in the reverse complementary orientation, i.e. as positive-strand sequences. The alignment was drawn in the Graphic View mode of the BioEdit program. For clarity, the primers and probe sequences were separated by tildes and numbered according to their M-segment positions. The dots indicate an identical nucleotide and the horizontal grey bar designates the threshold (≥0.5%). The shaded groups designates potentially critical M-segment variants. “Outgroup 1” contains sequences with at least one uncertainty (R, Y, N, etc), and “outgroup 2” encompasses an incomplete database of submissions. The “excluded” submissions contain no data.

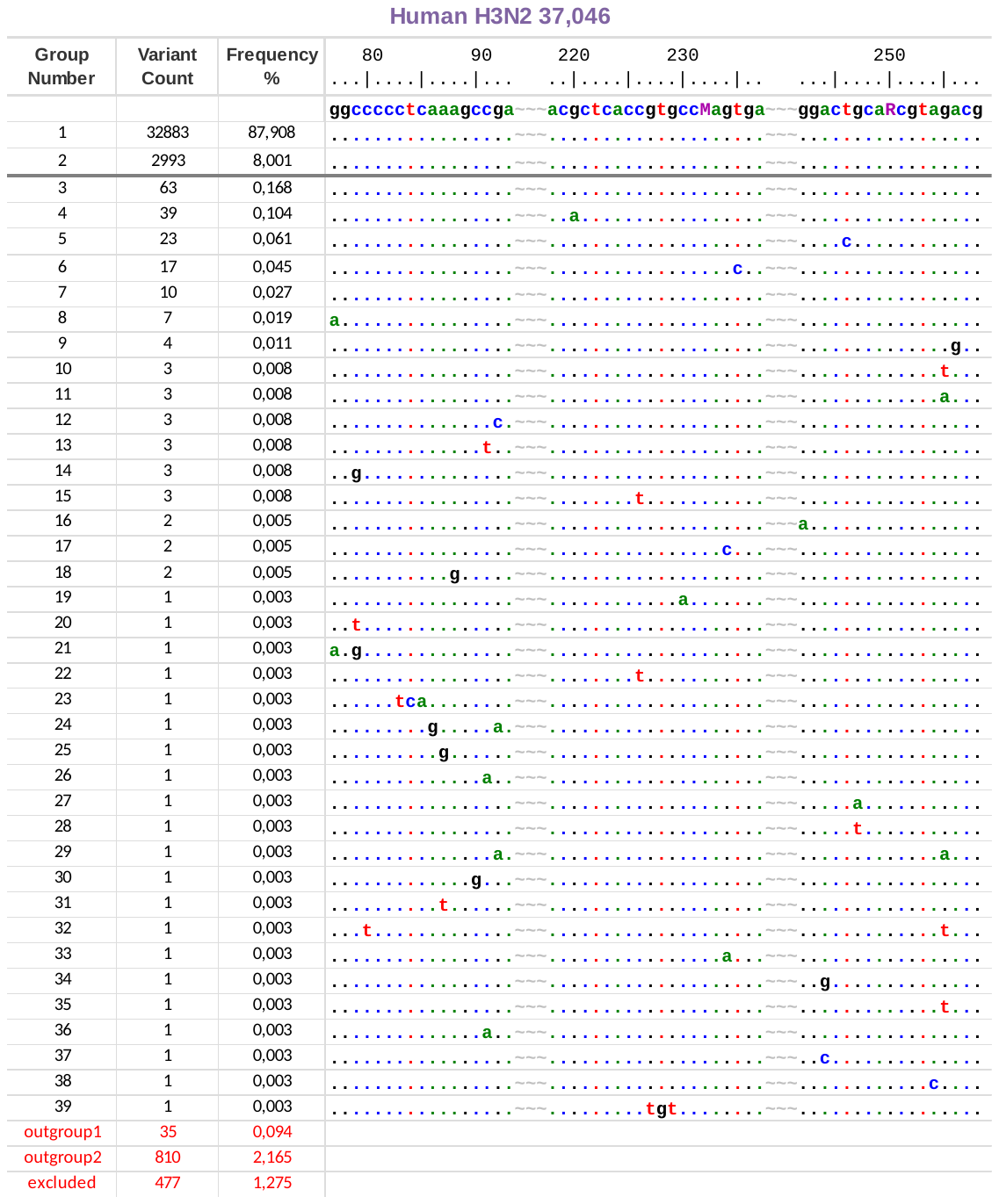
