## Supplementary material for "A universal RT-qPCR assay for “One Health” detection of influenza A viruses": Validation of the SVIP-MPv2 at APHA-Weybridge (UK) using avian, swine and human IAV isolates and clinical specimens from AIV- and SwIV-infected birds

**Supporting Information 3:** Validation of the SVIP-MPv2 at APHA-Weybridge (UK) using avian, swine and human IAV isolates and clinical specimens from AIV- and SwIV-infected birds and pigs respectively. References specific to this Supporting File are listed at the foot of this document, while any abbreviations which do not feature in the main text are also defined in this document.

**Methods:**

Avian viral isolates were obtained and propagated in embryonated fowls eggs (EFEs) by classical means and included AIVs (OIE, 2015a), and included AIVs (n=77), avian orthoavulavirses type-1 (AOAvV-1s, formerly known as avian paramyxoviruses type-1 (APMV-1s) which include the pigeon paramyxoviruses type-1; n=15) and infectious bronchitis virses (IBVs) (n=6). Swine influenza viruses (SwIVs; n=54) and six human IAVs were propagated similarly in EFEs or *in vitro* in MDCK cell culture (OIE, 2015b). Viral RNA was extracted manually from all virus isolates and the supernatant fluids from swine respiratory organ homogenates (n=7) (Slomka *et al.*, 2009 and 2010), while robotic extraction was employed to extract avian swab fluids (Slomka *et al.*, 2009). All animal experiments were approved by the local APHA Animal Welfare and Ethical Review Body to comply with the relevant European and UK legislation (Slomka *et al*., 2018). Isolate-derived viral RNA was tested as either neat or diluted (as indicated below), while clinical-specimen derived RNA (n=79) was tested undiluted. The following RT-qPCRs featured in the validation presented in this Supporting Information:

- **Three M-gene RT-qPCRs for generic AIV detection**:
  1. Spackman *et al.* (2002), applied as described by Slomka *et al.* (2009)
  2. SVIP-MPv1: M-gene RT-qPCR version 1: Nagy *et al*., 2010; applied as described by Slomka *et al*. (2018)
  3. SVIP-MPv2: M-gene RT-qPCR version 2 (Nagy *et al.*, this study)
- **Generic SwIV RT-qPCR**: M-gene ‘perfect match’ RRT-PCR ( Slomka *et al.* (2010a)
- **L-gene (MGB-2) RT-qPCR** for detection of (AOAvV-1s, formerly avian paramyxovirus type 1) viruses: Sutton *et al.* (2019).

All nucleic acid extracts were simultaneously tested using the above RRT-PCRs, as relevant to the below tabulated viral sample groups. All amplifications were carried out on Mx3005P qPCR System PCR thermocyclers (Agilent). The QuantiFast Probe RT-PCR Master Mix kit (Qiagen) was used in the SVIP-MPv2 assay at APHA-Weybridge (UK). Cq values obtained from all relevant tests are listed below in the final three columns of each table, with Cq 36 serving as the positive cut-off, based on previous IAV RT-qPCR validation work at APHA and outbreak experience (Slomka *et al*., 2009, 2010a and b).

**Results**

**Table A.** RNA extracted from AIV (EFE-grown isolates n=77) tested by the three M-gene RT-qPCRs, namely Spackman et al (2002), SVIP-MPv1 (Nagy et al., 2010) and SVIP-MPv2 (this study). Viral RNA was tested as either neat (undiluted) or diluted as indicated, with Cq values tabulated. All H5 and H7 isolates are LPAIVs unless indicated otherwise. #: A/goose/Guangdong/1/1996 (Gs/Gd) lineage H5Nx HPAIVs (clade in parenthesis).

| **ID** | **AIV subtype** | **AIV isolate name** | **Dilution of AIV RNA** | **M-gene (Spackman *et al* 2002)** | **M-gene (SVIP-MPv1) - QuantiFast kit** | **M-gene (SVIP-MPv2,this study) - QuantiFast kit** |
| --- | --- | --- | --- | --- | --- | --- |
| 1 | H1N1 | A/duck/Alberta/35/1976 | 1/100 | 23.68 | 24.59 | 25.12 |
| 2 | H2N3 | A/duck/Germany/1215/1973 | 1/100 | 23.1 | 25.94 | 26.38 |
| 3 | H3N2 | A/turkey/England/1969 | 1/100 | 29.37 | 30.3 | 31.53 |
| 4 | H4N6 | A/duck/Czechoslovakia/1956 | Neat | 17.22 | No Cq | 19.59 |
| 5 | H4N6 | A/chicken/England/476-016250/2014 | 1/1000 | 24.94 | 24.06 | 25.72 |
| 6 | H5N1 HP | A/chicken/Scotland/1959 | 1/100 | 25.09 | 27.42 | 27.89 |
| 7 | H5N1 HP | A/turkey/Turkey/1/2005 (clade 2.2) ^#^ | 1/100 | 28.27 | 27.74 | 28.15 |
| 8 | H5N1 HP | A/turkey/England/2614/2007 (clade 2.2) ^#^ | 1/100 | 25.33 | 23.76 | 24.62 |
| 9 | H5N1 HP | A/goose/Hungary/3413/2007 (clade 2.2) ^#^ | 1/100 | 22.2 | 17.7 | 18.84 |
| 10 | H5N2 | A/ostrich/Denmark/72420/1996 | 1/100 | 27.05 | 26.48 | 26.83 |
| 11 | H5N2 | A/mallard/Denmark/2006 | 1/100 | 19.32 | 19.09 | 20.03 |
| 12 | H5N2 | A/chicken/Italy/11VIR/7548/2012 | 1/100 | 26.02 | 17.2 | 18.63 |
| 13 | H5N3 | A/teal/ England/7394-2805/2006 | 1/100 | 23.72 | 24.68 | 25.32 |
| 14 | H5N5 HP | A/mute Swan/Croatia/102/16 (clade 2.3.4.4) ^#^ | 1/100 | 24.82 | 25.92 | 26.57 |
| 15 | H5N6 | A/duck/Potsdam/2216/1984 | 1/100 | 24.57 | 27.15 | 27.62 |
| 16 | H5N6 HP | A/mandarin duck/Korea/T102-1/2016 (clade 2.3.4.4c) ^#^ | 1/100 | 30.85 | 20.89 | 23.14 |
| 17 | H5N6 HP | A/broiler duck/Korea/H15/2016 (clade 2.3.4.4c) ^#^ | 1/1000 | 33.52 | 23.32 | 24.8 |
| 18 | H5N6 HP | A/chicken/Korea/HN1/2016 (clade 2.3.4.4c) ^#^ | 1/10000 | 35.28 | 26.43 | 27.99 |
| 19 | H5N6 HP | A/broiler duck/Korea/ES2/2016 (clade 2.3.4.4c) ^#^ | 1/10000 | 38.75 | 31.36 | 32.03 |
| 20 | H5N6 HP | A/spot-billed duck/Korea/WB141/2017 (clade 2.3.4.4c) ^#^ | 1/1000 | 33.65 | 24.06 | 25.83 |
| 21 | H5N6 HP | A/chicken/Greece/39/2017 (clade 2.3.4.4b) ^#^ | 1/10000 | 33.01 | 30.81 | 33.18 |
| 22 | H5N6 HP | A/mute swan/England/AVP-18-001986/2017 (clade 2.3.4.4b) ^#^ | 1/100 | 34.19 | 26.9 | 29.46 |
| 23 | H5N7 | A/duck/Denmark/64650/2003 | 1/100 | 26.95 | 30.16 | 30.69 |
| 24 | H5N8 HP | A/turkey/Ireland/1744/1983 | 1/100 | 24.37 | 24.1 | 24.79 |
| 25 | H5N8 HP | A/wigeon/Wales/52833/2016 (clade 2.3.4.4b) ^#^ | 1/1000 | 28.71 | 25.67 | 26.79 |
| 26 | H5N8 HP | A/domestic chicken/Bulgaria/295/2018 (clade 2.3.4.4b) ^#^ | Neat | 16.57 | 15.6 | 16.92 |
| 27 | H5N8 HP | A/commercial duck/Bulgaria/307/2018 (clade 2.3.4.4b) ^#^ | Neat | 15.33 | 14.93 | 16.67 |
| 28 | H5N8 HP | A/commercial duck/Bulgaria/309/2018 (clade 2.3.4.4b) ^#^ | Neat | 32.93 | 34.63 | 34.95 |
| 29 | H5N8 HP | A/domestic chicken/Bulgaria/333/2018 (clade 2.3.4.4b) ^#^ | Neat | 15.33 | 14.38 | 15.74 |
| 30 | H5N8 HP | A/domestic turkey/Bulgaria/336/2018 (clade 2.3.4.4b) ^#^ | Neat | 16.15 | 15.36 | 16.61 |
| 31 | H5N8 HP | A/pekin duck/Bulgaria/98/2/2018 (clade 2.3.4.4b) ^#^ | Neat | 19.59 | 18.66 | 19.83 |
| 32 | H5N9 | A/chicken/Italy/22A/1998 | 1/100 | 25.35 | 29.34 | 29.04 |
| 33 | H6N1 | A/turkey/England/198/2009 | 1/100 | 24.63 | 27.9 | 28.09 |
| 34 | H6N2 | A/turkey/Massachusetts/1965 | 1/100 | 23.47 | 25.95 | 25.37 |
| 35 | H7N1 | A/African starling/England/983/1979 | 1/100 | 26.02 | 30.05 | 30.39 |
| 36 | H7N1 HP | A/turkey/Italy/977/1999 | 1/100 | 24.58 | 23.91 | 25.86 |
| 37 | H7N1 | A/turkey/Italy/117/V00/2000 | 1/100 | 24.77 | 24.3 | 25.97 |
| 38 | H7N2 | A/chicken/Wales/1306/2007 | 1/100 | 24.25 | 25.55 | 25.74 |
| 39 | H7N3 | A/chicken/England/4054/2006 | 1/100 | 24.55 | 30.54 | 28.83 |
| 40 | H7N3 | A/chicken/Italy/11VIR/62/2012 | 1/100 | 23.13 | 22.13 | 23.69 |
| 41 | H7N7 | A/turkey/England/647/1977 | 1/100 | 25.54 | 25.17 | 24.93 |
| 42 | H7N7 | A/turkey/England/268/1996 | 1/100 | 26.06 | 29.42 | 29.41 |
| 43 | H7N7 | A/mallard/Sweden/100993/2008 | Neat | 22.48 | 21.53 | 24 |
| 44 | H7N7 HP | A/chicken/England/011406/2008 | 1:10000 | 28.86 | 27.21 | 28.53 |
| 45 | H7N7 | A/mallard duck/Italy/11VIR/540/2012 | 1/100 | 21.65 | 22.05 | 23.45 |
| 46 | H7N7 | A/chicken/Netherlands/15004744/2015 | Neat | 21.39 | 19.59 | 20.79 |
| 47 | H7N7 | A/mallard/Netherlands/19/2015 | Neat | 17.25 | 15.01 | 18.78 |
| 48 | H7N7 HP | A/chicken/England/26352/2015 | Neat | 17.24 | 15.43 | 18.12 |
| 49 | H7N9 | A/Anhui/1/2013 | 1/100 | 21.31 | 24.74 | 24.81 |
| 50 | H8N4 | A/turkey/Ontario/6118/1968 | 1/1000 | 30.27 | 25.76 | 27.97 |
| 51 | H9N2 | A/turkey/Wisconsin/1/1966 | 1/100 | 25.09 | 27.49 | 27.48 |
| 52 | H9N2 | A/pheasant/Republic of Ireland/PV18/1997 | Neat | 18.52 | 19.73 | 20.44 |
| 53 | H9N2 | A/chicken/Hong Kong/G9/97 | Neat | 15.5 | 14.92 | 15.71 |
| 54 | H9N2 | A/chicken/Germany/K1009/98/1998 | Neat | 17.97 | 16.9 | 17.9 |
| 55 | H9N2 | A/avian/Middle East/2/1998 | Neat | 15.44 | 15.31 | 16.22 |
| 56 | H9N2 | A/chicken/Republic of South Korea/99029/1999 | Neat | 19.14 | 18.62 | 20.8 |
| 57 | H9N9 | A/knot/England/SV497/2002 | 1/100 | 26.09 | 29.53 | 29.55 |
| 58 | H9N2 | A/chicken/India/4/2003 | Neat | 13 | 13.4 | 14.58 |
| 59 | H9N2 | A/chicken/Pakistan/47/2003 | Neat | 12.87 | 13.17 | 14.23 |
| 60 | H9N2 | A/shorebird/DE/9/2006 | Neat | 13.88 | 13.95 | 15.51 |
| 61 | H9N2 | A/duck/Hunan/1/2006 | Neat | 12.13 | 12.59 | 13.8 |
| 62 | H9N2 | A/mallard/England/6499/2006 | 1/100 | 22.46 | 21.51 | 22.71 |
| 63 | H9N2 | A/chicken/Bangladesh/301/2007 | Neat | 12.96 | 14 | 15 |
| 64 | H9N2 | A/mallard/Finland/Li13384/2010 | Neat | 18.04 | 20.1 | 21.98 |
| 65 | H9N2 | A/chicken/Iraq/30/2011 | Neat | 13.82 | 14.91 | 15.62 |
| 66 | H9N2 | A/turkey/England/2013 | 1/10000 | 25.09 | 24.4 | 25.74 |
| 67 | H9N2 | A/chicken/Nepal/2013 | 1/10000 | 26.17 | 25.28 | 26.45 |
| 68 | H10N4 | A/teal/Italy/11VIR/660/2012 | 1/100 | 21.32 | 20.62 | 22.42 |
| 69 | H10N7 | A/mallard/England/7495/2006 | 1/10 | 18.37 | 17.54 | 19.16 |
| 70 | H10N8 | A/mallard/Netherlands/06014516/2006 | 1/100 | 24.63 | 27.59 | 27.36 |
| 71 | H10N9 | A/RSA/Egyptian goose/238/1998 | 1/100 | 25.91 | 31.88 | 29.55 |
| 72 | H11N6 | A/duck/England/1956 | 1/100 | 24.18 | 30.25 | 28.49 |
| 73 | H12N5 | A/duck/Alberta/60/1976 | 1/100 | 22.59 | 26.08 | 24.92 |
| 74 | H13N6 | A/gull/Maryland/704/1977 | 1/10 | 23.09 | 26.08 | 28.41 |
| 75 | H14N6 | A/mallard/Gurjev/244/1982 | 1/1000 | 26.08 | 27.26 | 28.81 |
| 76 | H15N9 | A/wedge-tailed shearwater/Western Australia/2576/1979 | 1/100 | 25.73 | 29.31 | 29.55 |
| 77 | H16N3 | A/gull/Denmark/68110/2002 | 1/100 | 26.2 | 27.27 | 26.82 |

**Table B.** Cq values tabulated for swabs (oropharyngeal and cloacal; n=10) obtained from birds infected experimentally with two historic HPAIVs, RNA extracted and tested by the three relevant AIV M-gene RT-qPCRs.

| **ID** | **AIV subtype** | **Isolate name** | **Sample details** | **M-gene (Spackman *et al* 2002)** | **M-gene (SVIP-MPv1) - QuantiFast kit** | **M-gene (SVIP-MPv2,this study) - QuantiFast kit** |
| --- | --- | --- | --- | --- | --- | --- |
| 1 | H5N8 HP | A/turkey/Ireland/1744/1983 | Duck 155, 1dpi, cloacal swab | 33.98 | 32.01 | 32.98 |
| 2 | H5N8 HP | A/turkey/Ireland/1744/1983 | Duck 156, 1dpi, cloacal swab | 27.54 | 24.61 | 25.26 |
| 3 | H5N8 HP | A/turkey/Ireland/1744/1983 | Duck 159, 1dpi, cloacal swab | 29.41 | 26.91 | 27.78 |
| 4 | H5N8 HP | A/turkey/Ireland/1744/1983 | Duck 158, 1dpi, Buccal swab | 32.46 | 30.81 | 31.32 |
| 5 | H7N1 HP | A/ostrich/Italy/984/2000 | Chicken, oro-pharyngeal swab | No RNA available | 23.64 | 24.84 |
| 6 | H7N1 HP | A/chicken/Italy/1279/1999 (1/10 dilution) | Chicken, oro-pharyngeal swab | 34.82 | 33.3 | 34.08 |
| 7 | H7N1 HP | A/ostrich/Italy/984/2000 (1/10 dilution) | Chicken, oro-pharyngeal swab | 31.62 | 28.99 | 30.09 |
| 8 | H7N1 HP | A/ostrich/Italy/984/2000 (1/10 dilution) | Chicken, cloacal swab | 29.68 | 27.28 | 27.85 |
| 10 | H7N1 HP | A/ostrich/Italy/984/2000 (1/10 dilution) | Chicken, cloacal swab | 29.56 | 26.9 | 27.7 |

**Table C.** Testing of representative mammalian influenza A virus isolates (n=60) which included SwIV isolates (n=54) and six human influenza A isolates, latter indicated by *italic text*. Extracted viral RNA was tested by the three relevant M-gene RRT-PCRs, namely the ‘perfect match’ (Slomka *et al* 2010), SVIP-MPv1 (Nagy *et al* 2010) and SVIP-MPv2 (this study) assays, with Cq values tabulated. Other than the subtype, further details of the SwIVs’ origins are indicated where known: H1N2 = reassortant human-like H1N2; avH1N1 = avian-like H1N1 origin; H1N1 pdm 09 = evolved from the prototype H1N1 pandemic 2009 IAV; H1N2/avH1N1 = external glycoproteins of human-reassortment (H1N2) origin but six internal genes derived from avH1N1; H3N2 = external glycoproteins of human origin.

| **ID** | **Influenza A virus subtype** | **Isolate name** | **Dilution of extracted SwIV RNA** | **M-gene ‘perfect match’; Slomka et al 2010a)** | **M-gene (SVIP-MPv1) - QuantiFast kit** | **M-gene (SVIP-MPv2,this study) - QuantiFast kit** |
| --- | --- | --- | --- | --- | --- | --- |
| 1 | H1N2 | A/swine/England/028256/14 | 1/100 | 24.37 | 32.11 | 25.57 |
| 2 | avH1N1 | A/swine/England/033403/14 | 1/100 | 27.18 | 25.98 | 27.17 |
| 3 | H1N2 | A/swine/England/034079/14 | 1/100 | 23.68 | 24.05 | 27.92 |
| 4 | H1N2 | A/swine/England/087378/14 | 1/100 | 23.39 | 22.47 | 24.23 |
| 5 | H1N2 | A/swine/England/096724/14 | 1/100 | 25.48 | 24.51 | 26.46 |
| 6 | H1N2 | A/swine/England/097085/14 | 1/100 | 21.77 | 21.12 | 22.37 |
| 7 | H1N2 | A/swine/England/019733/14 | 1/100 | 26.76 | 25.65 | 27.17 |
| 8 | H1N2 | A/swine/England/105127/14 | 1/100 | 22.36 | 21.62 | 23.34 |
| 9 | H1N2 | A/swine/England/035708/15 | 1/100 | 22.41 | 21.45 | 23.32 |
| 10 | H1N2 | A/swine/England/132457/15 | 1/100 | 22.8 | 21.73 | 23.52 |
| 11 | H1N2 | A/swine/England/220198/15 | 1/100 | 20.71 | 19.57 | 21.23 |
| 12 | H1N2 | A/swine/England/042792/16 | 1/100 | 23.8 | 23.4 | 24.88 |
| 13 | avH1N1 | A/swine/England/044314/16 | 1/100 | 26.47 | 25.48 | 26.71 |
| 14 | H1N2 | A/swine/England/161262/16 | 1/100 | 21.71 | 21.07 | 22.58 |
| 15 | H1N2 | A/swine/England/174832/16 | 1/100 | 23.56 | 22.34 | 23.92 |
| 16 | H1N2 | A/swine/England/062058/18 | 1/100 | 22.15 | 21.74 | 23.47 |
| 17 | H1N2 | A/swine/England/062942/18 | 1/100 | 20.05 | 19.36 | 20.63 |
| 18 | H1N2 | A/swine/England/064519/18 | 1/100 | 21.18 | 20.98 | 23.26 |
| 19 | H1N2 | A/swine/England/208046/18 | 1/100 | 22.12 | 21.56 | 22.61 |
| 20 | avH1N1 | A/swine/England/267/2007 | 1/100 | 30.42 | 31.28 | 30.67 |
| 21 | H1N2/avH1N1 | A/swine/England/SP15001249/2011 | 1/100 | 27.47 | 26.41 | 27.85 |
| 22 | H1N2 | A/swine/England/33780/06 | 1/100 | 25.51 | 25.14 | 25.36 |
| 23 | H1N1 pdm 09 | A/swine/England/4202/2013 | 1/100 | 27.43 | 27 | 28.58 |
| 24 | H1N2 | A/swine/England/041118/2013 | 1/100 | 26.51 | 26.96 | 27.12 |
| *25* | *H1N1 Human* | *A/Solomon Islands/3/06* | *Neat* | *19.65* | *15.73* | *17.83* |
| *26* | *H1N1 pdm09 human* | *A/California/04/09* | *Neat* | *13.47* | *14.5* | *16.38* |
| *27* | *H1N1 pdm09 human* | *A/Michigan/45/2015* | *Neat* | *13.97* | *17.61* | *18.28* |
| *28* | *H1N1 pdm09 human* | *A/Israel/Q-504/2015* | *Neat* | *16.82* | *17.94* | *19.34* |
| 29 | avH1N1 | A/swine/Gent/13/17 | Neat | 15.76 | 17.99 | 18.7 |
| 30 | avH1N1 | A/swine/Gent/173/15 | Neat | 14.46 | 15.3 | 16.28 |
| 31 | avH1N1 | A/swine/Gent/138/17 | Neat | 15.1 | 15.79 | 16.85 |
| 32 | avH1N1 | A/swine/Gent/P20/18 | Neat | 18.01 | 13.86 | 15.08 |
| 33 | avH1N1 | A/swine/Gent/8/18 | Neat | 13.46 | 14.29 | 15.45 |
| 34 | avH1N1 | A/swine/Gent/61/14 | Neat | 15.38 | 14.72 | 16.18 |
| 35 | avH1N1 | A/swine/Gent/62/15 | Neat | 18.16 | 16.14 | 16.83 |
| 36 | avH1N1 | A/swine/Gent/157/17 | Neat | 18.89 | 15.49 | 16.86 |
| 37 | avH1N1 | A/swine/Gent/31/18 | Neat | 18.49 | 14.48 | 15.11 |
| 38 | avH1N1 | A/swine/Gent/32/18 | Neat | 14.11 | 15.24 | 16.41 |
| 39 | H1N1 | A/swine/Spain/40250-1/2016 | Neat | 17.45 | 16.36 | 16.46 |
| 40 | H1N1 | A/swine/Spain/40250-2/2016 | Neat | 13.48 | 17.51 | 17.94 |
| 41 | H1N1 | A/swine/Spain/45690-2/2017 | Neat | No Ct | 14.84 | 15.48 |
| 42 | H1N1 | A/swine/Spain/40340-1/2017 | Neat | 12.12 | 14.82 | 16.05 |
| 43 | H1N1 | A/swine/NL/Rhezerveen-CV19121A/2012 | Neat | 17.6 | 15.76 | 16.3 |
| 44 | H1N2 | A/swine/Gent/5/17 | Neat | 13.49 | 14.92 | 15.26 |
| 45 | H1N2 | A/swine/Gent/P18/18 | Neat | 18.61 | 14.08 | 15.51 |
| 46 | H1N2 | A/swine/Gent/7/18 | Neat | 14.54 | 13.99 | 14.9 |
| 47 | H1N2 | A/swine/Spain/45700-1/2016 | Neat | 18.8 | 15.1 | 15.3 |
| 48 | H1N2 | A/swine/Spain/40340-2/2017 | Neat | 13.9 | 17.18 | 18.45 |
| 49 | H1N2 | A/swine/Spain/45600-1/2017 | Neat | 19.8 | 14.88 | 15.9 |
| 50 | H1N2 | A/swine/Spain/45600-2/2017 | Neat | 19.09 | 14.58 | 16.46 |
| 51 | H1N2 | A/swine/Spain/45600-3/2017 | Neat | 19.25 | 14.24 | 15.43 |
| 52 | H1N2 | A/swine/Spain/45600-4/2017 | Neat | 19.61 | 14.94 | 15.04 |
| 53 | H1N2 | A/swine/Scotland/410440/94 | Neat | 18.28 | 17.08 | 17.9 |
| *54* | *H3N2 human* | *A/Victoria/361/11* | *Neat* | *20.15* | *15.07* | *16.18* |
| *55* | *H3N2 human* | *A/Texas/50/2012* | *Neat* | *21.99* | *17.29* | *18.32* |
| *56* | H3N2 | A/swine/Gent/48/17 | Neat | 16.96 | 14.33 | 15.61 |
| 57 | H3N2 | A/swine/Gent/17/14 | Neat | 16.53 | 15.08 | 15.72 |
| 58 | H3N2 | A/swine/Gent/27/14 | Neat | 15.02 | 15.52 | 15.88 |
| 59 | H3N2 | A/swine/Gent/280/13 | Neat | 14.76 | 16.48 | 16.28 |
| 60 | H3N2 | A/swine/Spain/45690-1/2016 | Neat | No Ct | 17.46 | 17.37 |

**Table D.** Testing of total RNA extracts from respiratory tissues of SwIV-infected pigs (n=7) from the field in the UK. Testing was done using the three relevant M-gene RT-qPCRs, namely the ‘perfect match’ (Slomka *et al*., 2010a), SVIP-MPv1 (Nagy *et al*., 2010) and SVIP-MPv2 (this study) assays, with Cq values tabulated. All six internal genes of the seven infecting SwIVs were of H1N1 pandemic 2009 IAV, but HA and neuraminidase gene origins reflected reaasortment events, as indicated: pdm 09 = Both glycoprotein and the six internal gene segments evolved from the prototype H1N1 pandemic 2009 IAV (i.e. un-reassorted genomes); N2 – hu = N2 is neuraminidase of human-like origin; H1N2 - hu = Both glycoprotein genes are of non-pandemic 2009 human origin.

| **ID** | **SwIV subtype** | **Clinical specimen ID, i.e. SwIV isolate name** | **M-gene ‘perfect match’; Slomka et al 2010a)** | **M-gene (SVIP-MPv1) - QuantiFast kit** | **M-gene (SVIP-MPv2,this study) - QuantiFast kit** |
| --- | --- | --- | --- | --- | --- |
| 1 | H1N1 pdm 09 | A/swine/England/SA15-059903/2018 | 24.44 | 24.57 | 25.89 |
| 2 | H1N2, N2 - hu | A/swine/England/SA14-204123/2018 | 28.52 | 29.02 | 28.97 |
| 3 | H1N2 - hu | A/swine/England/SA15-058962/2018 | 22.3 | 22.9 | 23.15 |
| 4 | H1N2 - hu | A/swine/England/SA24-031878/2013 | 30.4 | 32.53 | 33.81 |
| 5 | H1N2 - hu | A/swine/England/SA14-155224/2016 | 26.65 | 27.9 | 29.4 |
| 6 | H1N1 pdm 09 | A/swine/England/SA15-943261/2016 | 30.75 | 33.13 | 32.91 |
| 7 | H1N1 pdm 09 | A/swine/England/014408/2016 | 30.61 | 33.44 | 33.46 |

**Table E.** RNA extracted from avian orthoavulavirus type 1 (AOAvV-1, formerly known as avian paramyxovirus type 1 [APMV-1]) isolates (n=15) used to validate the diagnostic specificity of the SVIP-MPv1 (Nagy et al., 2010) and SVIP-MPv2 (this study) M-gene RT-qPCRs, with the L-gene (MGB-2) RT-qPCR (specific for AOAvV-1s, Sutton et al., 2019) Cq values also shown. The list includes two pigeon paramyxovirus type 1 (PPMV-1) isolates. RNA extracted from all 15 egg-grown AOAvV-1 isolates were tested undiluted (neat).

| **ID** | **AOAvV-1 isolate name** | **AOAvV-1 lineage** | **L-gene (MGB-2) –Quantifast kit; Sutton et al (2019)** | **M-gene (SVIP-MPv1) - QuantiFast kit** | **M-gene (SVIP-MPv2,this study) - QuantiFast kit** |
| --- | --- | --- | --- | --- | --- |
| 1 | AOAvV-1/chicken/Australia/Queensland/V4/1966 | Lineage 1 | 9.58 | No Cq | No Cq |
| 2 | AOAvV-1/chicken/Northern Ireland/Ulster/2C | Lineage 1 | 29.7 | No Cq | No Cq |
| 3 | AOAvV-1/chicken/USA/Hitchner B1/1947 | Lineage 2 | 8.93 | No Cq | No Cq |
| 4 | AOAvV-1/chicken/USA/LaSota/1946 | Lineage 2 | 9.09 | No Cq | No Cq |
| 5 | AOAvV-1/chicken/India/Mukteswar/1945 | Lineage 3a | 10.03 | No Cq | No Cq |
| 6 | AOAvV /chicken/UK/Herts33/1933 | Lineage 3b | 17.8 | No Cq | No Cq |
| 7 | AOAvV-1/chicken/USA/211472/2003 | Lineage 3c | 10.96 | No Cq | No Cq |
| 8 | AOAvV-1/chicken/Iraq/AG68/1968 | Lineage 4a | 11.33 | No Cq | No Cq |
| 9 | AOAvV-1/partridge/UK/7575/2006 | Lineage 4b | 10.13 | No Cq | No Cq |
| 10 | PPMV-1/pigeon/UK/015874/2015 | Lineage 4b | 11.35 | No Cq | No Cq |
| 11 | PPMV-1/pigeon/UK/GB1168/1984 | Lineage 4b | 10.65 | No Cq | No Cq |
| 12 | AOAvV-1/chicken/Cyprus/AV000632/2013 | Lineage 5a | 9.55 | No Cq | No Cq |
| 13 | AOAvV-1/chicken/Nepal/8-43/2010 | Lineage 5b | 11.89 | No Cq | No Cq |
| 14 | AOAvV-1/teal/Finland/Li3111-2008/2013 | Lineage 6 | 20.43 | No Cq | No Cq |
| 15 | AOAvV-1/chicken/Ireland/IECK90187/1990 | Lineage 6 | 22.14 | No Cq | No Cq |

**Table F.** Six infectious bronchitis virus (IBV) isolates used to validate the diagnostic specificity of the Spackman et al (2002) and the SVIP-MPv1 (Nagy et al., 2010) and SVIP-MPv2 (this study) assays. IBV RNA was tested undiluted, with no Cq values registered by any of the three M-gene RT-qPCRs.

| **ID** | **IBV isolate name** | **IBV sample details** | **M-gene (Spackman *et al* 2002)** | **M-gene (SVIP-MPv1) - QuantiFast kit** | **M-gene (SVIP-MPv2,this study) - QuantiFast kit** |
| --- | --- | --- | --- | --- | --- |
| 1 | IBV Qx strain | Neat egg fluid | No Cq | No Cq | No Cq |
| 2 | IBV strain 793/B | Neat egg fluid | No Cq | No Cq | No Cq |
| 3 | IBV strain M41 | Neat egg fluid | No Cq | No Cq | No Cq |
| 4 | IBV 755/03 (CP) | Neat egg fluid | No Cq | No Cq | No Cq |
| 5 | IBV strain D1466 | Neat egg fluid | No Cq | No Cq | No Cq |
| 6 | IBV strain D274 | Neat egg fluid | No Cq | No Cq | No Cq |

**Graphical representation of swab testing (n=69) from an AIV outbreak or AIV experimentally-infected chickens**

Four regression graphs (A, B, C and D; below) illustrating a comparison of Cq values obtained by two M-gene RT-qPCRs: SVIP-MPv1 (Nagy et al., 2010) and SVIP-MPv2 (this study) from a total of 69 chicken swabs obtained from a H7N7 HPAIV (2015, UK) outbreak in layer hens (A; n=39, mix of oropharyngeal and cloacal) and from experimental infections with three other AIV subtypes (B, C and D; n=30). In B, specific pathogen free (SPF) chickens were infected as per the statutory intravenous pathogenicity index (IVPI) test (OIE 2015). In C and D, SPF chickens were infected with 6 log_10_EID_50_ via the ocular-nasal route at 6 and 3 weeks age respectively. Cq 36 positive cut-off is indicated by the broken lines. Regression graphs were assembled by Prism (Graphpad) software:

**A:**

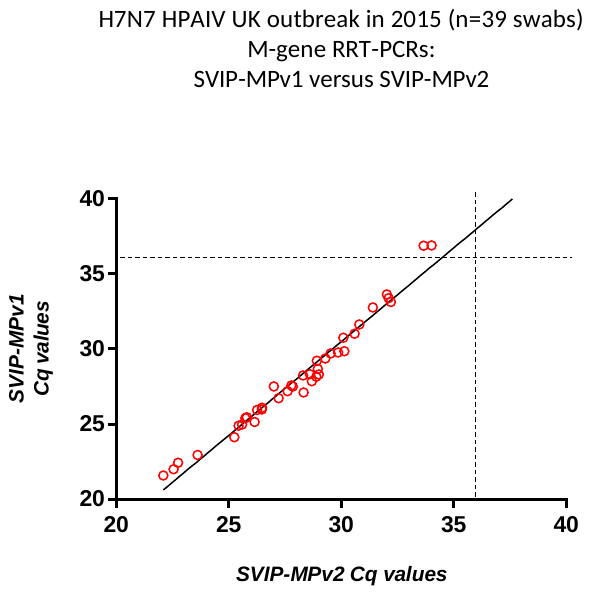

**B:**
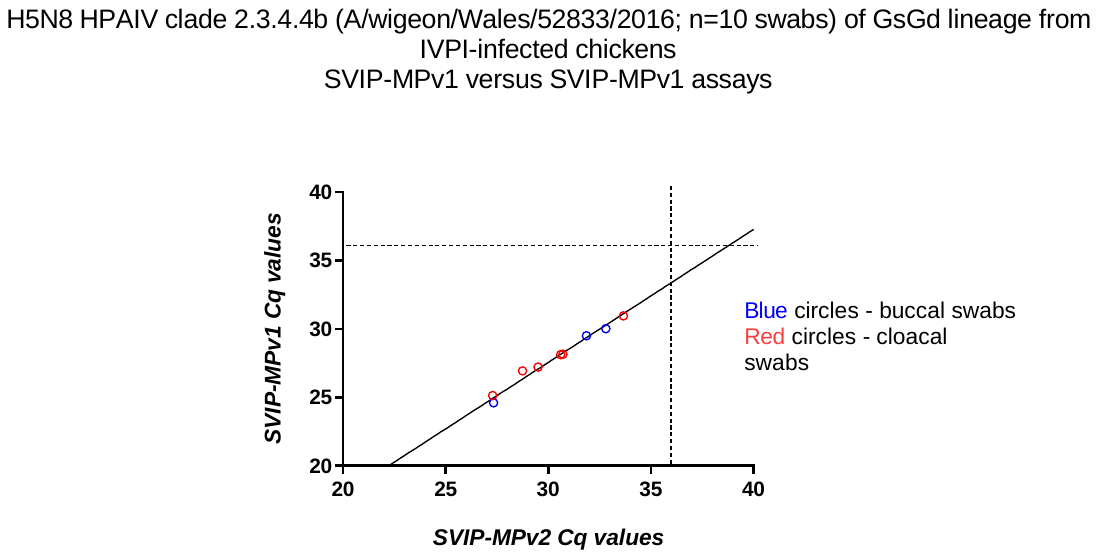

**C:**

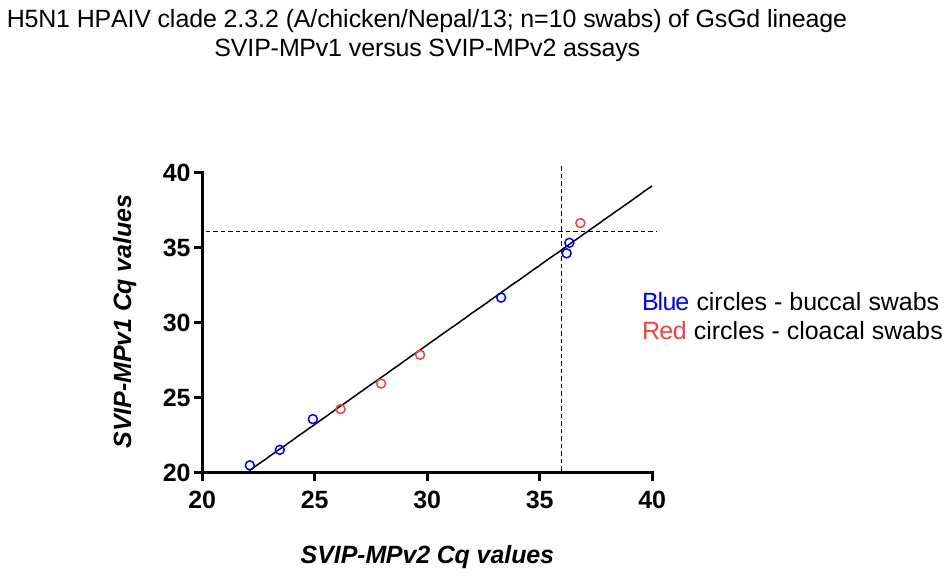

**D:**

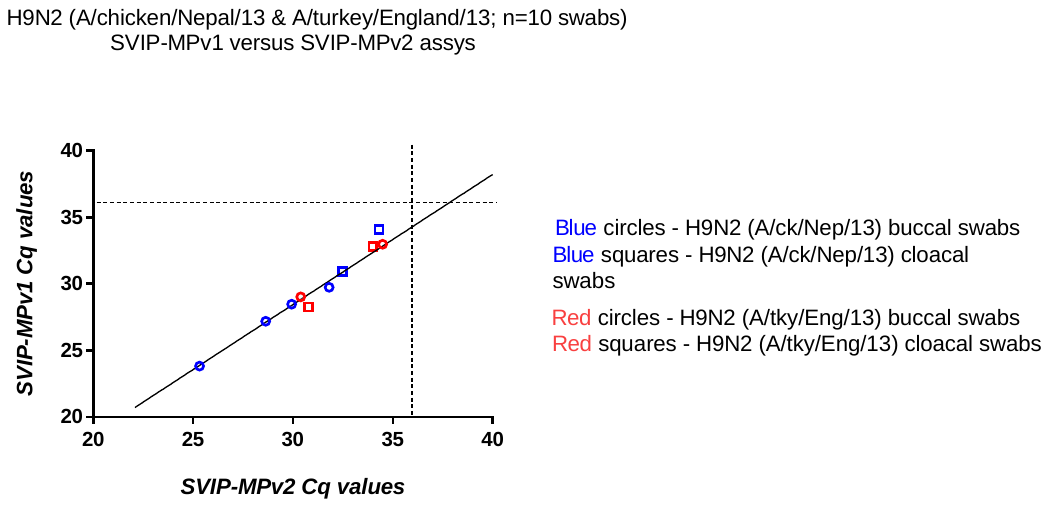
